# Quantification method affects replicability of eQTL analysis, colocalization, and TWAS

**DOI:** 10.1101/2025.08.20.671303

**Authors:** S. Taylor Head, Sean T. Bresnahan, Yung-Han Chang, Vihan Shah, Nolan Cole, William Wu, Arjun Bhattacharya

## Abstract

Expression quantitative trait loci (eQTL) mapping, colocalization, and transcriptome-wide association studies (TWAS) are widely used to contextualize GWAS loci, yet the impact of upstream RNA-seq processing choices on these analyses remains critically unexplored. Two poorly characterized decisions include the transcriptomic reference annotation used for quantification of short-read RNA-sequencing samples, which defines transcript composition and structure, and the quantification method itself. Through a comprehensive analysis spanning 48 tissues in GTEx and seven widely studied and diverse phenotypes, here we find that these choices substantially affect eQTL detection and gene expression prediction with significant downstream impact on colocalization and TWAS results. Concordant results across pipelines are further enriched for longer genes, genes with more isoforms, and greater expression heritability. Our findings demonstrate that seemingly minor methodological decisions substantially affect these common analyses, highlighting the need for standardized practices to ensure reproducible genetic association studies.

## INTRODUCTION

Contextualizing variant-to-trait associations identified through genome-wide association studies (GWAS) often relies on expression quantitative trait loci (eQTL) mapping, colocalization and transcriptome-wide association studies (TWAS)^1,2^. Reference eQTL datasets for these integrative analyses are often based on short-read RNA-sequencing (RNA-seq) data. Two critical decisions in short-read RNA-seq analysis workflows involve (1) the choice of transcriptomic annotation that defines gene, exon, and transcript boundaries for expression quantification and (2) the method itself for quantification. GENCODE annotations are regularly updated to incorporate newly discovered transcripts and refined gene models^3^. Quantification methods also vary considerably, from alignment using STAR^4^ followed by featureCounts, htseq-count, or RSEM^5–7^ to alignment-free methods such as Salmon and kallisto^8,9^, each with distinct assumptions about transcript structure and quantification accuracy.

Short-read RNA-seq-based eQTL colocalization and TWAS continue to be commonly used to identify mechanisms underlying GWAS-identified loci. As of June 2025, we found 124 pre-printed or published studies that included gene-level eQTL colocalization or TWAS with short-read RNA-seq indexed in PubMed Central (**Supplemental Table S1**). Of these studies, over 90% quantify expression through an alignment-based approach. These studies also used a variety of transcriptomic annotations ranging from GENCODE v17 to GENCODE v38, with a few using *de novo* transcriptomes or combining datasets quantified based on multiple annotations. Many of these studies cite the original publication performing the sequencing for RNA-seq processing details, which is the minimal practice when leveraging established resources. Nonetheless, only 10 of these studies directly report the quantification method or reference transcriptome either in the main text or supplemental methods. This highlights that these parameters are not yet routinely considered key methodological details needing explicit documentation.

While previous studies have compared quantification methods on simulated data^10,11^, their relative performance for genetic association analyses using real human tissue data remains unclear. Here, we extracted raw RNA-seq FASTQ files from 48 tissues from the Genotype-Tissue Expression Project (GTEx v8, **Methods**). We quantified gene-level expression using GENCODE v27, v38, v45, and Ensembl v115 transcript annotations through either STAR alignment with featureCounts^4,6^ or RSEM^7^, or pseudo-alignment with Salmon^8^ or kallisto^9^. We then conducted eQTL analysis using QTLtools^1^, eCAVIAR colocalization with large-scale GWAS for seven phenotypically diverse traits (bipolar disorder^12^, body mass index^13^, breast cancer^14^, height^14,15^, prostate cancer^16^, schizophrenia^17^, and type 2 diabetes^18^), and TWAS^2^ (**Supplemental Table S2, Methods**).

## RESULTS

Across the four annotations considered, Ensembl is the most comprehensive and catalogs more than half a million individual transcripts. Within the GENCODE releases, the number of genes remained relatively similar, but the number of annotated transcripts increased with time to approximately 250,000 in the v45 release (**Supplemental Figure S1**). We find significant discordance in genes with significant eQTLs (eGenes) and those with predicted expression at R^2^ > 0.01, both across quantification method and across annotation. Across 48 tissues, eGene calls are ∼39-43% discordant across method and ∼32-39% discordant across annotation (**Figure 1A-B**). This discordance holds regardless of the p-value threshold used to define eGenes (**Supplemental Figure S2**). Additionally, changing splice junction overhang parameters in STAR had minimal to no impact on QTL discovery. To assess whether these discrepancies generalize beyond GTEx, we performed an additional analysis in an independent dataset of 150 healthy breast tissue samples and a subset of quantification methods and annotations. We see similar discordance rates across annotation and greater discordance by quantification method compared to GTEx breast tissue samples and two other size-matched tissues in GTEx (**Supplemental Figure S3**).

**Figure 1.**
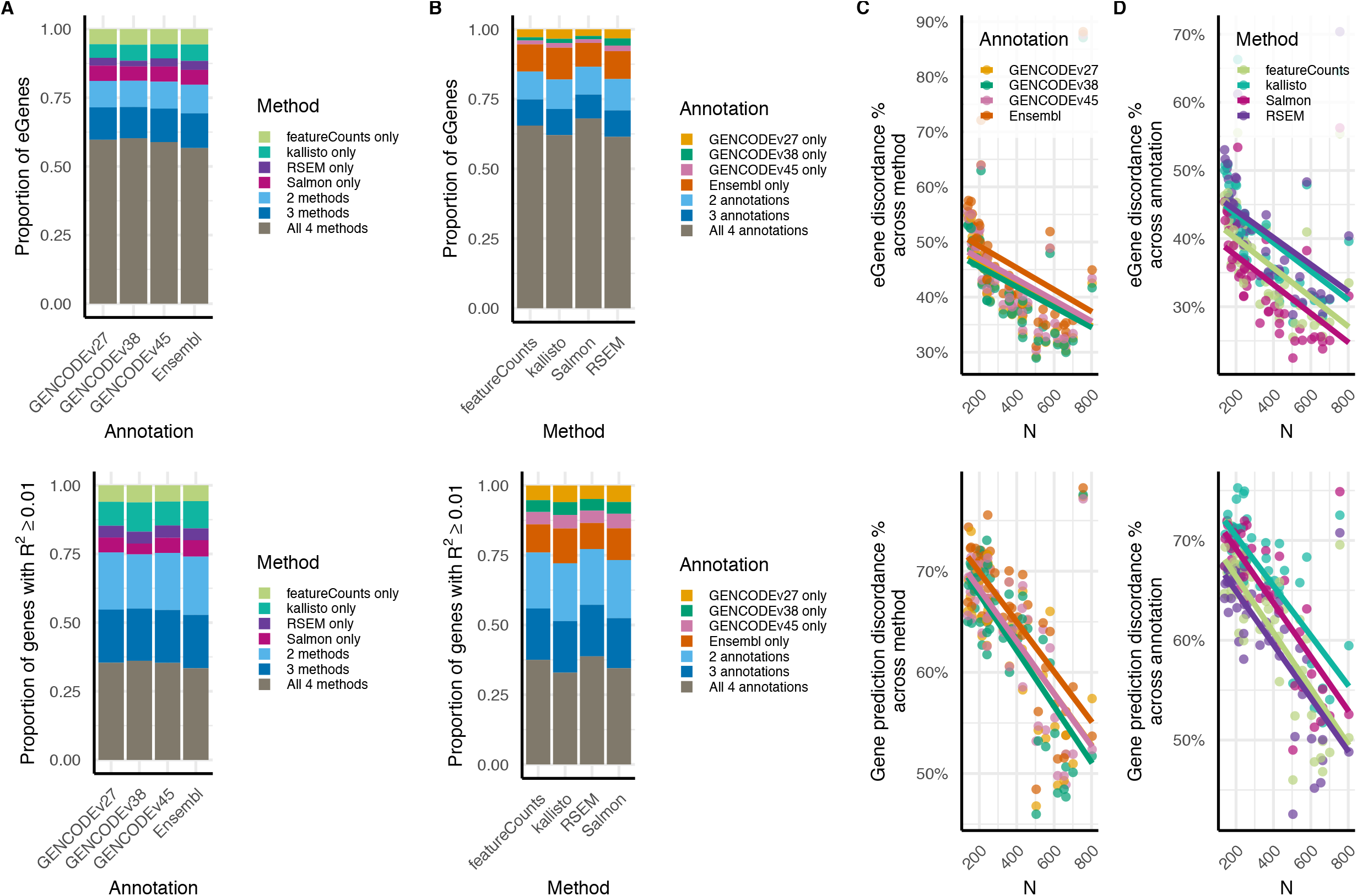
Quantification method and transcriptomic annotation affect eGene identification and gene expression prediction. **(A)** Proportion of eGenes (top) and genes with R^2^ > 0.01 (bottom) identified by each combination of quantification methods across transcriptome annotations (x-axis). **(B)** Proportion of eGenes (top) and predicted genes (bottom) identified by one or more annotations by quantification method (x-axis). **(C)** Method discordance percentage vs. tissue sample size for eGene calling (top) and expression prediction (bottom). **(D)** Annotation discordance percentage vs. tissue sample size for eGene calling (top) and expression prediction (bottom).

We find eQTL signal strength is greatest among eGenes concordant across both axes, followed by those concordant across quantification methods alone (**Supplemental Figure S4A**). Within a tissue, eGene set overlap and correlation in lead eQTL effect sizes are highest among runs sharing the same method and lowest among runs differing in both method and annotation (**Supplemental Figure S4B-C**). We also find that the number of discordant eGenes reflects chromosome length and relative abundance of gene biotypes (**Supplemental Figure S5-7**). The majority of annotation- and method-concordant eGenes are protein-coding, whereas the majority of annotation- and method-specific eGenes are lncRNAs and pseudogenes. Even so, across tissues, nearly 10,000 discordant eGenes specific to only one method or one annotation are protein-coding. Annotation-concordant eGenes robustly detected across all annotations (**Supplemental Figure S8-10**) and method-concordant eGenes robustly detected across methods (**Supplemental Figures S11-13**) tend to have more transcripts, contain more exons, and are correspondingly longer than discordant eGenes. Additionally, eGenes identified consistently across all annotations (**Supplemental Figure S14**) and across all methods (**Supplemental Figure S15**) are enriched for genes supported by independent disease evidence from Open Targets after excluding RNA expression-based evidence (e.g., clinical, genetic, somatic mutations, animal models). This suggests that reproducibly detected regulatory targets may preferentially implicate biologically relevant genes with orthogonal support for disease association. However, when restricting the analysis to only protein-coding eGenes to control for potential bias of Open Target scores to favor well-annotated genes, the enrichment differential diminished. Here, there are often comparable proportions of high-scoring Open Targets genes across concordance classes (**Supplemental Figures S16-17**). Thus, phenotypically relevant genes with orthogonal disease evidence may be frequently missed as a direct consequence of quantification method and transcript annotation choice.

Multi-SNP predictions of gene expression exhibit far more discordance: ∼64-67% discordance across method and ∼61-67% discordance across annotation (**Figure 1A-B**), owing potentially to uncertainty in variable selection during penalized regression. In fact, the number of SNPs included in expression prediction models for the same gene in the same tissue varies greatly, when comparing across method and annotation (**Supplemental Figures S18-S19**), which leads to differences in interpretation of the polygenicity of the gene’s expression. These discordance rates generally decrease with increasing sample size, yet a non-negligible portion of discordance remains even in the tissues with the largest sample sizes (**Figure 1C-D, Supplemental Tables S3-6, Supplemental Figures S20-27**), suggesting that these discrepancies may not be fully mitigated even in large datasets.

At these sample sizes, these discrepancies in QTL calling and expression prediction have stark effects on downstream colocalization and TWAS. Colocalization analysis across seven diverse GWAS phenotypes reveals substantial discordance in gene identification across both quantification methods and reference annotations. While the independent GWAS loci tagged by colocalized genes show discordance of ∼55-63% (**Figure 2A**), the specific genes identified through colocalization at these loci exhibit more marked discordance. Annotation-based comparisons between transcriptomes show less than 30% concordance in colocalized gene identification, with the remainder showing some degree of annotation-specificity (**Figure 2B**). Even more pronounced discordance is observed across quantification methods, where up to 79% of colocalized genes are identified in three or fewer methods, and we see the majority of colocalized genes are method-specific rather than consistently identified in more than one approach (**Figure 2B, Supplemental Figures S28-31, Supplemental Tables S7-10**). These results reflect findings from previous studies demonstrating that annotation and methodological choices substantially impact RNA-seq quantification accuracy^19,20^.

**Figure 2:**
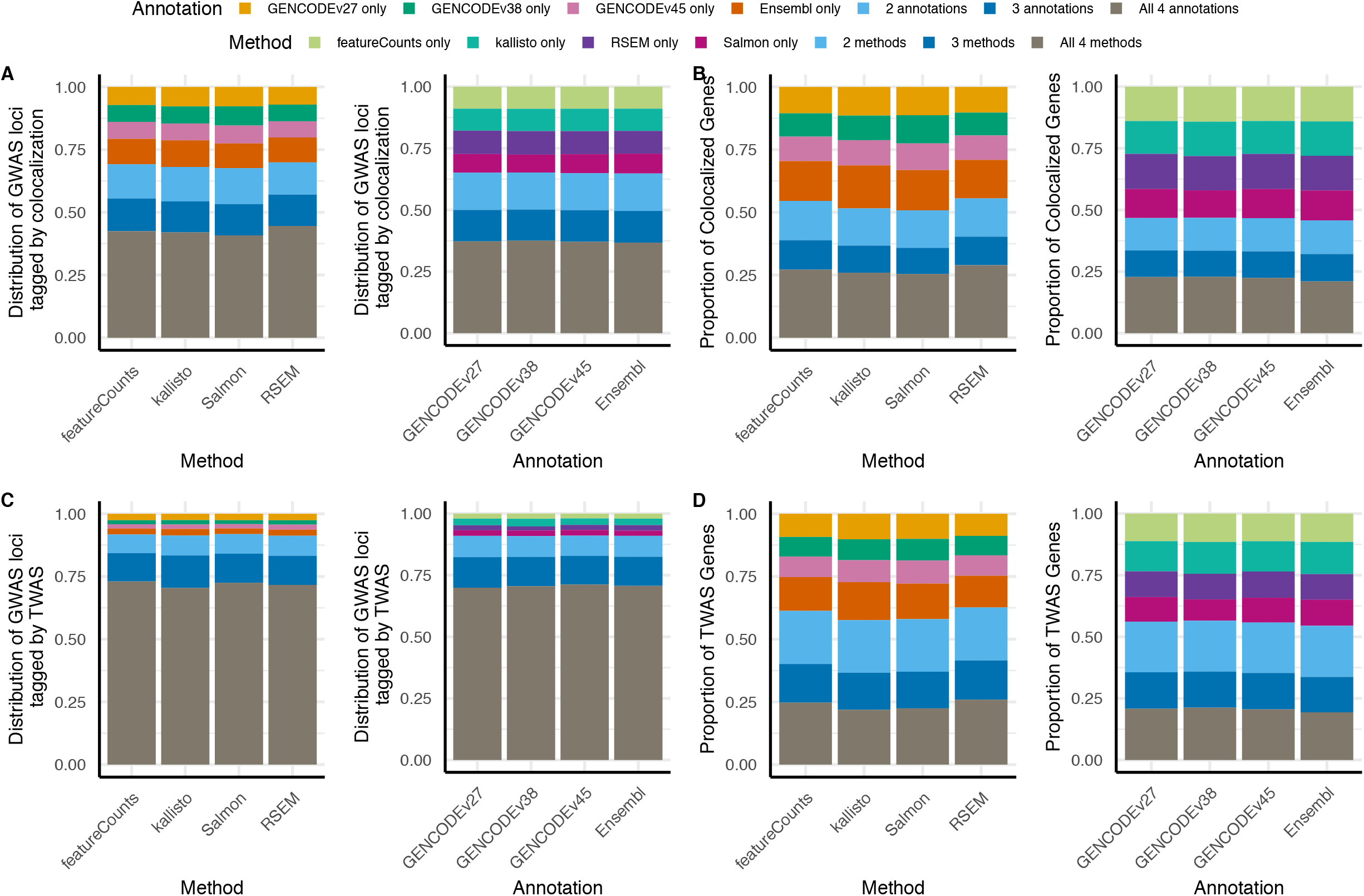
Differences in quantification method and transcriptomic annotation lead to discordances in GWAS colocalization and TWAS. Across 48 GTEx tissues, proportion (Y-axis) of all GWAS loci tagged through colocalization **(A)** or TWAS **(C)** using QTL data quantified with different methods or using different reference annotations. Across 48 GTEx tissues, proportion (Y-axis) of all colocalized (B) or TWAS-prioritized (D) genes within 1 Mb of a GWAS locus across quantification method and reference annotation.

In contrast to the colocalization results, across both quantification method and annotation, the independent GWAS loci that are tagged by TWAS-identified genes within 1 Mb are generally concordant, with ∼70-73% concordance (**Figure 2C, Supplemental Figures S32-33, Supplemental Tables S11-14**). However, TWAS mimics the strong discrepancies we observed in colocalization between tagged GWAS loci and the actual genes being prioritized at these loci. The specific genes identified at these loci across method and across annotation are quite different and show up to 80% and 78% discordance across method and annotation, respectively (**Figure 2D, Supplemental Figures S34-35**). TWAS identifies a distinct set of genes per locus, depending on quantification method and annotation (**Supplemental Figure S36**). Although this discrepancy could imply that GWAS variants potentially affect transcription of multiple genes at the locus, we believe, rather, it raises a concern for replicability, especially since the underlying GWAS and eQTL datasets are the same. As a cautionary note, TWAS associations do not imply shared underlying variants, and a strong eQTL or GWAS effect can drive a significant TWAS signal even under weak LD between the two^21^. As a result, different annotations or quantification methods may flag different genes at the same locus simply because each captures a different LD-tagging eQTL as its lead signal. This highlights the complementary value of colocalization for evaluating candidate genes at a locus.

To test whether pleiotropic eQTL regulation could account for discordant colocalization and TWAS results, we examined whether the lead eQTL for each gene was shared with other genes in the same tissue, but no consistent pattern was observed across colocalization/TWAS discordance groups (**Supplemental Figure S37-S38**). Further, method- and annotation-discordant colocalization and TWAS results are not solely attributable to discordant eGenes in the majority of cases, with 42-83% of downstream discordant genes classified as fully concordant eGenes across both annotation and method (**Supplemental Figure S39**). Among fully concordant eGenes, downstream concordance in colocalization and TWAS was observed at rates of 26-39% and 31-64%, respectively, across the seven phenotypes examined (**Supplemental Figure S40**). These findings indicate that considerable pipeline-level variability in colocalization and TWAS results arises at steps downstream of eGene identification and that eGene concordance alone does not guarantee concordant disease-gene prioritization.

To more directly evaluate factors in genetic architecture that may drive such discordance while accounting for realistic methodological and technical considerations, we developed a simulation framework directly modeling true underlying gene expression and simulating the corresponding short-read RNA-seq FASTQ files from it (**Methods**). Using this framework, we specifically examined the impact of two quantification methods and three GENCODE annotations on eGene identification in N = 500 samples. This approach allowed us to assess whether concordant and discordant eGenes were systematically associated with particular eQTL or sequencing characteristics. Although we observed minor differences in the effect of quality control steps like TPM filtering on the retention of heritable genes prior to QTL analysis by method and annotation (**Supplemental Figure S41**), we find expression heritability strongly associated with concordance. eGenes detected consistently across quantification methods or annotations exhibited substantially higher expression heritability than method- or annotation-specific eGenes (**Figure 3**). This finding is consistent with the expectation that genes with stronger genetically regulated components of expression are more robust to technical perturbations introduced by different processing pipelines. In contrast, genes with weaker genetic effects lie closer to significance thresholds. This makes their detection more sensitive to differences in read assignment and expression estimation. We also observed a modest tendency for concordant eGenes to have a greater number of annotated isoforms.

**Figure 3:**
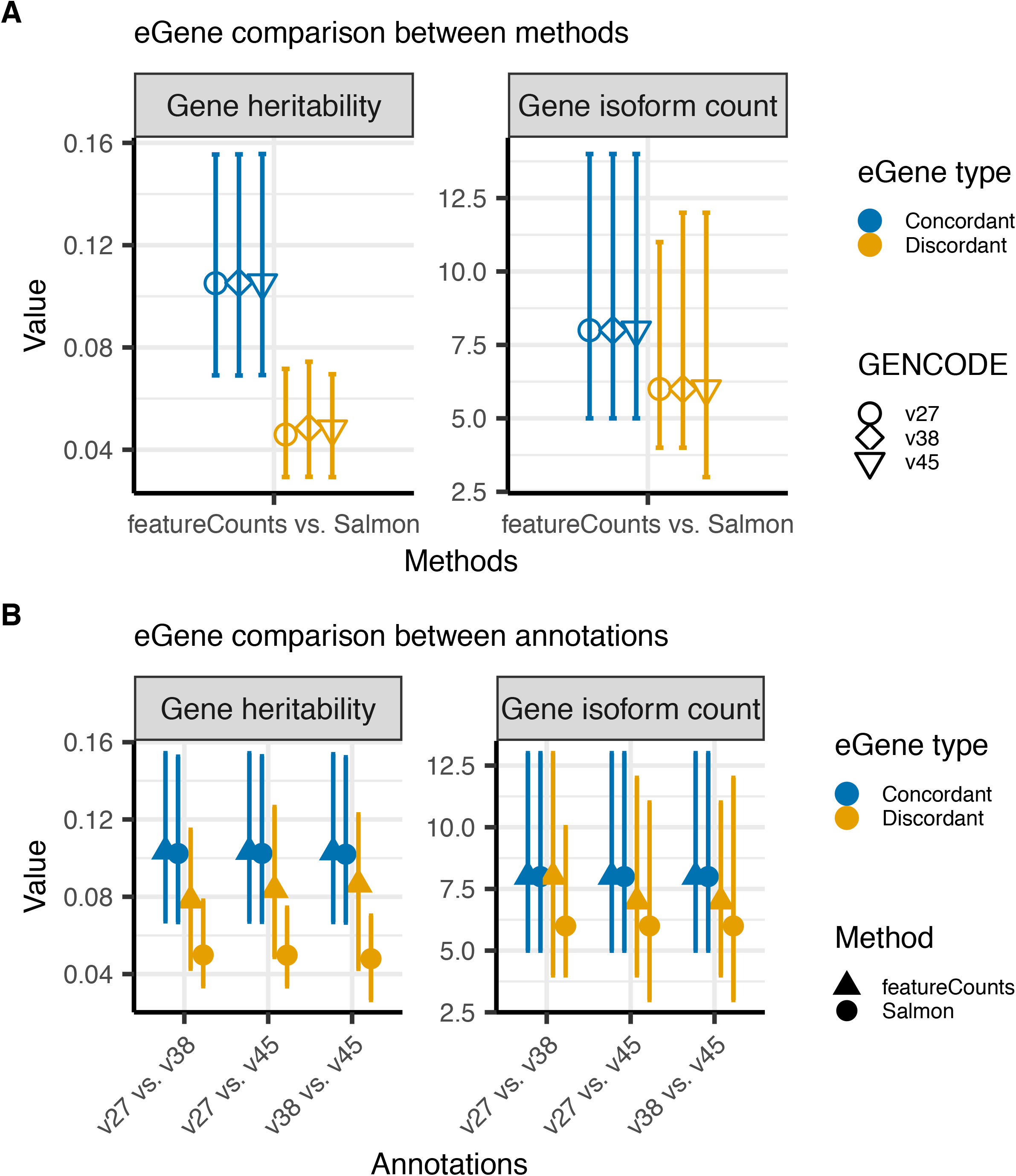
Differences in gene heritability and isoform count by discordant versus concordant eGenes in simulation analysis. Expression was simulated for autosomal protein-coding genes in GENCODE v38 using genotypes from N = 500 randomly selected European-ancestry 1000 Genomes Phase 3 individuals. We simulated short-read RNA-seq data and quantified under each setting (Salmon or featureCounts; GENCODE v27, v38 or v45). We used these data to compare the properties of eGenes consistently identified across annotations and quantification settings (concordant) versus those detected in only a subset of settings (discordant). Points indicate median values and error bars denote the interquartile range (25^th^-75th percentiles). Results correspond to 9,993 simulated eGenes detected in at least one setting. (**A**) Comparison across quantification methods while holding GENCODE annotation constant. (**B**) Comparison across GENCODE annotations while holding quantification method constant. Concordant eGenes exhibited higher expression heritability and a modest tendency toward greater annotated isoform counts than discordant eGenes.

## DISCUSSION

Our analysis shows that seemingly minor methodological choices in quantification method or transcript annotation substantially affect QTL results. These effects are notable even at the gene level and likely will be more pronounced for transcript- and exon-level analyses. As recent QTL studies are increasingly focusing on isoform expression and splicing, annotation choice becomes critical: larger and newer annotation versions introduce more isoforms and refined exon boundaries that can dramatically shift quantification and association outcomes. Additionally, as annotations vary by tissue and cell type, these choices have major implications for studying context-specific regulation. Our results show that two representative methodological discrepancies (transcript annotation and quantification method) diminish with increasing sample size and highlight the value of larger molecular phenotype datasets, although not enough to entirely remove all discordance. Stemming from our simulation findings, genes reproducibly detected across analytical approaches tended to have higher expression heritability, suggesting that increased sample sizes primarily improve concordance by enabling more robust detection of genes with weaker genetically regulated components of expression. Recent work further reinforces this need, showing that robust QTL-GWAS colocalization may require far greater sample sizes than currently available^22^. Together, these findings support the need for a strong commitment to expanding transcriptomic datasets. Nevertheless, even in the largest datasets, a non-negligible degree of discordance remained, indicating that increased power alone cannot fully overcome technical differences between processing pipelines: short-read data lack the resolution to capture complex and tissue-specific transcript structures. To address this, long-read sequencing must play a complementary role in future eQTL studies, enabling more accurate quantification of nuanced, context-specific transcript structures.

Another important implication of our findings is that no single transcript annotation can currently be considered universally optimal. Although GENCODE remains a widely used reference and provides broad coverage across tissues and cell types, emerging tissue-specific and long-read-derived annotations may better capture biologically relevant transcript diversity in particular cellular contexts. In recent work, our group performed both eGene and TWAS/colocalization analysis for breast cancer risk with *de novo* long-read transcriptomes and combined annotations integrating established GENCODE transcripts with novel long-read isoforms^23^. We identified regulatory signals that would have been missed using any single reference alone. Consequently, the choice of annotation should be guided by the goals of the analysis and the biological system under investigation rather than by convention. Annotation-free approaches to short-read RNA-seq quantification, which do not rely on a predefined transcript catalog, represent a promising alternative^24–27^. We therefore advocate that new annotations and quantification strategies, including annotation-free methods, be evaluated through rigorous benchmarking against existing approaches and underscore explicit reporting of these choices and their motivation. Future work will be needed to determine how annotation-free approaches compare with standard and tissue-specific long-read annotations in terms of replicability in eQTL mapping and downstream findings.

We conclude by advocating the field of functional genomics to adopt stronger methodological standards to ensure the reproducibility, transparency, and relevance of eQTL and TWAS analyses. First, rigorous reporting practices are imperative. Details about important processing choices like quantification methods and the transcriptomic annotations used should be treated as foundational, on par with reporting genome builds, to enable accurate interpretation and cross-study comparisons. Encouragingly, efforts such as the eQTL Catalogue^28^ and FIVEx^29^ have made important strides toward the harmonized processing of eQTL data across studies and tissues. These projects demonstrate both the feasibility and value of standardized pipelines. However, such efforts have not yet been universally adopted, and key parameters potentially underlying discovery discordance in many published analyses remain underreported.

Second, the community must establish frameworks for regularly updating foundational resources. Widely used pre-computed datasets such as GTEx eQTL maps and TWAS models from FUSION or PredictDB^30^ are often treated as static references, despite rapid improvements in gene annotations and analytic methods. To reflect the current state of genomic knowledge and avoid the propagation of outdated findings, we advocate for community-supported systems to re-quantify and refresh these key resources as methods and annotations evolve. As complex deep learning methods become increasingly central to transcriptome analysis^31^, the need for dynamic model updating will become even more imperative, ensuring that these models are trained on the most accurate and up-to-date representations of transcriptomic complexity. We consider these challenges presented here as a methodological opportunity to ensure reproducibility of computational inference from transcriptomic datasets.

## ACKNOWLEDGEMENTS

We thank Michael Love and Sara Lindström for discussion.

## AUTHOR CONTRIBUTIONS STATEMENT

A.B. conceived and supervised the study and wrote the original draft of the manuscript. S.T.H., S.T.B., and A.B. performed all analyses and substantially revised the manuscript. Y.-H.C., V.S., N.C., and W.W. contributed to literature searches, data processing, and analyses. All authors reviewed and contributed to the writing and editing of the final manuscript.

## METHODS

### Genotype processing

SNP genotype calls from GTEx v8^32^ were derived from Whole Genome Sequencing data, filtering out SNPs with missingness >5%, minor allele frequency (MAF) less than 1%, or that deviated from HWE at *P*□< □10^−6^. In the supplemental analysis of 150 independent African-ancestry healthy breast tissue samples from Ping et al.^33^, we obtained genotype data generated from the Illumina MEGA platform and previously imputed with TOPMed. We identified six sample pairs with estimated KING kinship coefficients exceeding 0.0884 and retained 144 unrelated samples for further analysis. We filtered out variants with MAF <1%, those deviating from HWE at P < 10^−6^, and those with missingness exceeding 5%. Genotype data were processed with PLINK v2.00a6LM^34^ and tabix v0.2.6^35^,

## RNA-seq processing

We quantified distinct GTEx v8^32^ RNA-seq samples for 48 tissues using four commonly used tools. For all, we initially derived raw FASTQ files from the source BAMs by coordinate-sorting with SAMtools v1.16.1^36^ and converting to paired-end FASTQ using `samtools fastq`.

First, we estimated transcript abundance using Salmon v1.10.2^8^ in mapping-based mode. We built Salmon indices for decoy-aware transcriptomes for GENCODE v27, GENCODE v38, GENCODE v45, and Ensembl v115 transcript sequences and the full GRCh38 reference genome as decoy sequences^3^. We ran Salmon on paired-end FASTQ files with mapping validation and correction for sequencing bias. We imported transcript-level abundance estimates into R using tximeta v1.22.1^37^ and summarized to the gene level using `summarizeToGene`.

Second, we estimated transcript abundance using an alternative pseudoalignment-based method, Kallisto v0.48.0^9^, with annotation-specific transcriptome indices and default paired-end quantification. We again imported transcript-level abundance estimates with tximeta^37^ and summarized to the gene level using `summarizeToGene`.

Third, we performed read alignment using STAR v2.7.4a^4^, with a two-pass-like approach optimized for spliced transcript detection. The alignment was configured for high sensitivity to splice junctions (`--alignSJoverhangMin 8`, `--alignSJDBoverhangMin 1`) and permissive mismatch thresholds (`--outFilterMismatchNmax 999`, `--outFilterMismatchNoverReadLmax 0.04`). We allowed up to 20 multimapping reads per alignment and enabled long intron detection up to 1 Mb. Output was generated as sorted BAM files with comprehensive alignment attributes. We then generated gene-level counts using featureCounts v2.0.6^6^ and assigning paired-end fragments to annotated exon features and summing counts across exons belonging to the same gene.

Lastly, we estimated transcript abundance using RSEM v1.3.3^7^ with annotation-specific reference indices on paired-end FASTQ files using STAR as the underlying aligner and assuming unstranded libraries. We imported abundance estimates into R using tximeta and summarized to the gene level using `summarizeToGene`.

After each quantification procedure, we used edgeR v4.0.16 for TMM-normalization of gene-level counts. Before downstream analysis, we retained genes with TMM-normalized counts per million (CPM) > 0.1 in at least 25% of samples. We then log2-transformed CPM values (after adding a pseudocount of one) and residualized expression by all tissue-specific covariates (clinical, demographic, genotype principal components (PCs), and expression PEER factors) used in the original QTL analyses in GTEx. To ensure that comparisons are fair across method and annotation, we only retained genes that were not filtered in all datasets (across method and across annotation).

For the independent panel of breast tissue samples from Ping et al.^33^, we restricted quantification to two methods (Salmon, featureCounts) and transcriptome annotations to GENCODE, as illustrative, with both pipelines as described above. We residualized gene expression for top 15 genetic PCs and age.

### eQTL mapping and colocalization

We conducted eQTL analysis using QTLtools v1.3.1 with rank normalization of gene expression^1^. P-value correction was conducted using the Beta-distribution approximation using 1000 permutations. This test uses linear regression and assumes an approximately linear genotype-phenotype relationship and appropriately modeled residuals. For eGenes with adjusted P < 0.05 that were within 1 Mb of a GWAS SNP with P < 5e-8, we then conducted colocalization using eCAVIAR v2.2, assuming up to 2 causal variants per gene^38^. Prior to colocalization, we first harmonized all GWAS summary statistics. We independently munged each dataset using MungeSumstats v1.10.1^39^ with an imputation quality filter of INFO ≥ 0.3 and removed strand-ambiguous variants (A/T and C/G SNPs). All other parameters were left at default settings.

### Transcriptome-wide association study

We followed the standard FUSION^2^ pipeline for predicting gene expression models using the most predictive in cross-validation of elastic net^40^, best linear unbiased predictor from a linear mixed model^41^, or Sum of Single Effects regression^42^. We then incorporated these models GWAS summary statistics with a weighted-burden test, running trait mapping only for genes predicted at cross-validation adjusted R^2^ > 0.01. We used in-sample linkage disequilibrium from GTEx to compute the weighted burden test statistic. We aggregated P-values from multiple tissue-specific gene associations using the aggregated Cauchy association test and considered P < 2.5 × 10^−6^ as transcriptome-wide significant^43^. We define an independent GWAS locus by identifying lead SNPs with minimum P < 5×10^−8^ in a 1 Mb window^44^.

### Simulation study

We randomly selected 500 European-ancestry samples from 1000 Genomes Phase 3^45^ to serve as our simulation study sample. We restricted consideration to autosomal protein-coding genes present in the GENCODE v38 annotation. For each gene, we simulated a true underlying isoform expression matrix **Y** (500 x k isoforms) = XB+E where **X** is the genotype matrix of *w* isoform-level QTLs (isoQTLs) within a 250kb *cis* window across the *k* isoforms. **B** is the w×k matrix of isoQTL effect sizes. We assumed c=4 causal isoQTLs with MAF > 0.05 per transcript isoform and assumed 50%, 75%, or 100% of these eQTLs were shared among all isoforms of a given gene. We drew per-gene heritability (h^2^) from Beta(2, 40). We simulated effect sizes for shared isoQTLs from MVN(**0**, Σ_**B**_) with covariance off-diagonal elements from Uniform(0.7, 0.9). We drew isoform-specific eQTL effects independently from 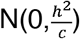. Causal SNPs were selected at random from the *cis*-window, and genotypes were mean-centered and unit-scaled. Residual covariance Σ_**E**_ had off-diagonal elements from Uniform(0, 0.5). We projected Σ_**B**_ and Σ_**E**_ to the nearest positive definite matrix as needed. We defined gene-level expression as the row sum of **Y**. Simulated heritability distribution is shown in **Supplemental Figure S42**. We then exponentiated simulated expression values to enforce non-negative means and passed to `simReads()` from Rsubread v2.16.1^46^ to generate paired-end reads (library size 30M, read length 75 bp).

We compared expression quantification and subsequent eGene discovery with two methods (Salmon and featureCounts) and three annotations (GENCODE v27, v38, v45). For each method, we performed isoform- and gene-level expression quantification as described above (**RNA-seq processing**). We retained only those genes with TPM >0.1 in at least 25% of samples and TMM-normalized expression. We performed eQTL analysis using QTLtools v1.3.1 with rank normalization of gene expression^1^ and identified eGenes with adjusted P < 0.05. P-value adjustment was conducted using the Beta-distribution approximation using 1000 permutations.

## DATA AVAILABILITY

GTEx genotype and short-read RNA-seq data were downloaded from dbGaP phs000424.v10.p2. Independent genotype and short-read RNA-seq data of healthy breast tissue from Ping et al.^33^ were downloaded from dbGap phs003535.v1.p1. 1000 Genomes genotype data for simulation samples were downloaded from https://www.internationalgenome.org/data-portal/sample. Available GWAS summary statistics were accessed from the GWAS Catalog with accession numbers listed in **Supplemental Table S2**. We provide nominal eQTL summary statistics for all *cis*-SNPs across each of the three GENCODE annotation versions examined on Zenodo (10.5281/zenodo.20765828)^47^. Results are available for all 48 GTEx tissues and include both Salmon and STAR+featureCounts results as representative quantification methods.

## CODE AVAILABILITY

All scripts for this analysis are available at github.com/bhattacharya-a-bt/QuantMethod_QTL_Comparison (10.5281/zenodo.23070543)^48^.

