## Supplemental Figures S1-S42 for "Quantification method affects replicability of eQTL analysis, colocalization, and TWAS"

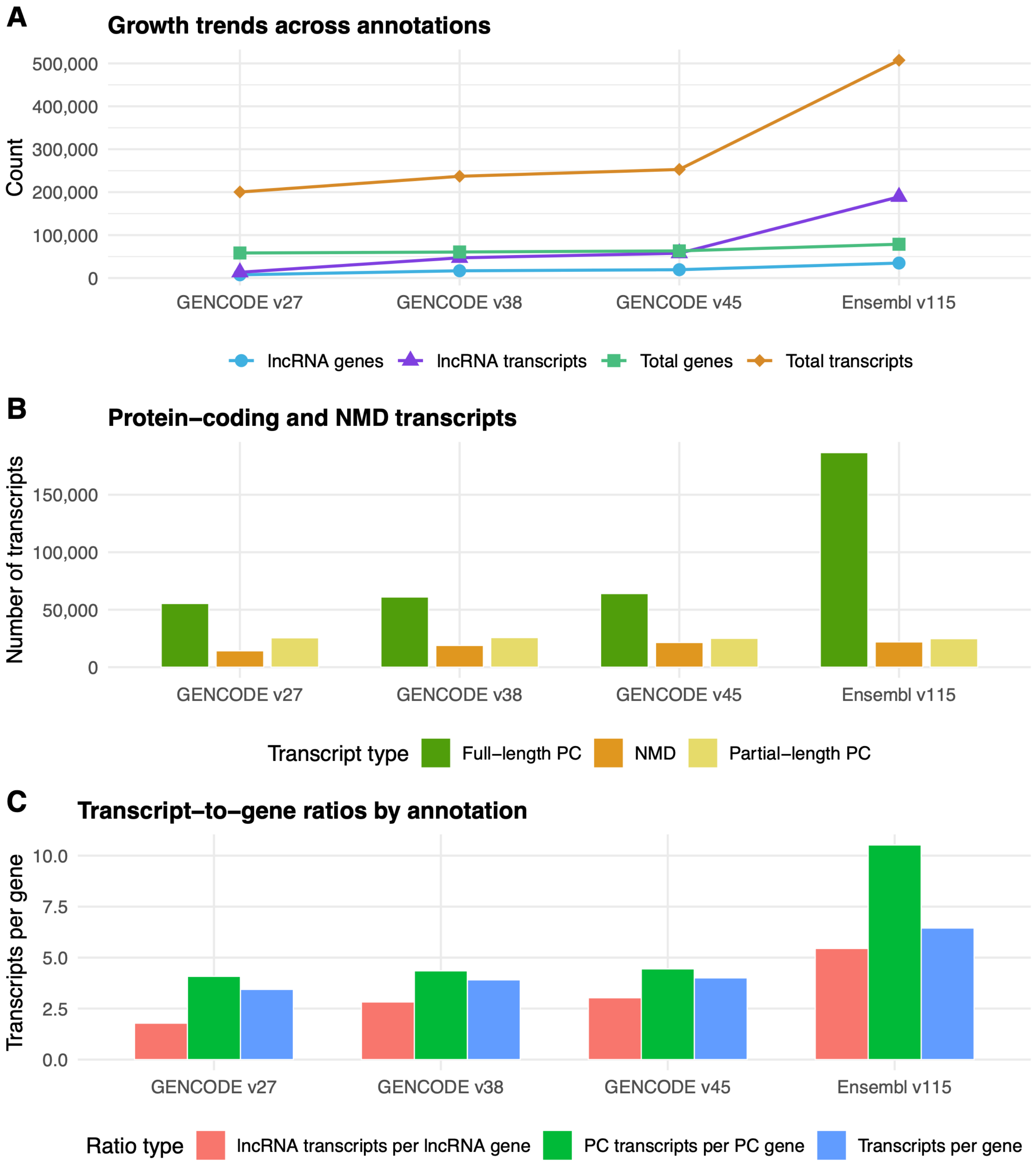


**Figure S1: (A)** Comparison of number of non-coding RNAs and total genes by transcriptome annotation. **(B)** Comparison of transcript types (full-length protein-coding, nonsense mediated decay, and partial length protein-coding) across annotations. **(C)** Ratio of transcripts to genes across annotations, stratified by non-coding RNAs and protein-coding (PC) genes.

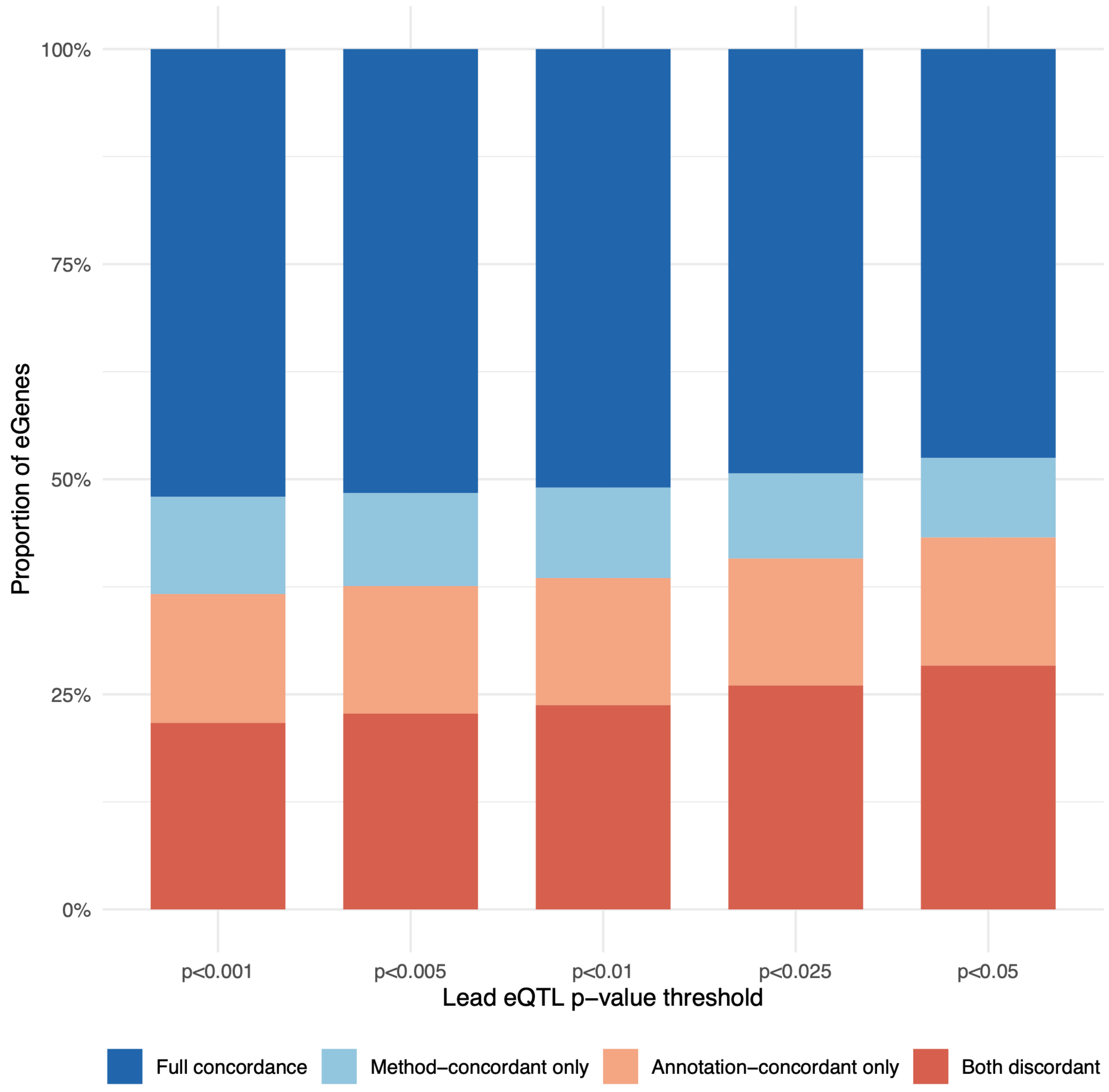


**Figure S2.** For a given adjusted p-value threshold (x-axis), we show the proportion of eGenes that are concordant across quantification methods (i.e., identified as eGenes by each method), concordant across transcriptome annotations, concordant across both methods and annotations (full concordance, dark blue), or discordant across both methods and annotations (both discordant, red). The p-value threshold corresponds to the p-value correction conducted using the Beta-distribution approximation using 1000 permutations to define eGenes.

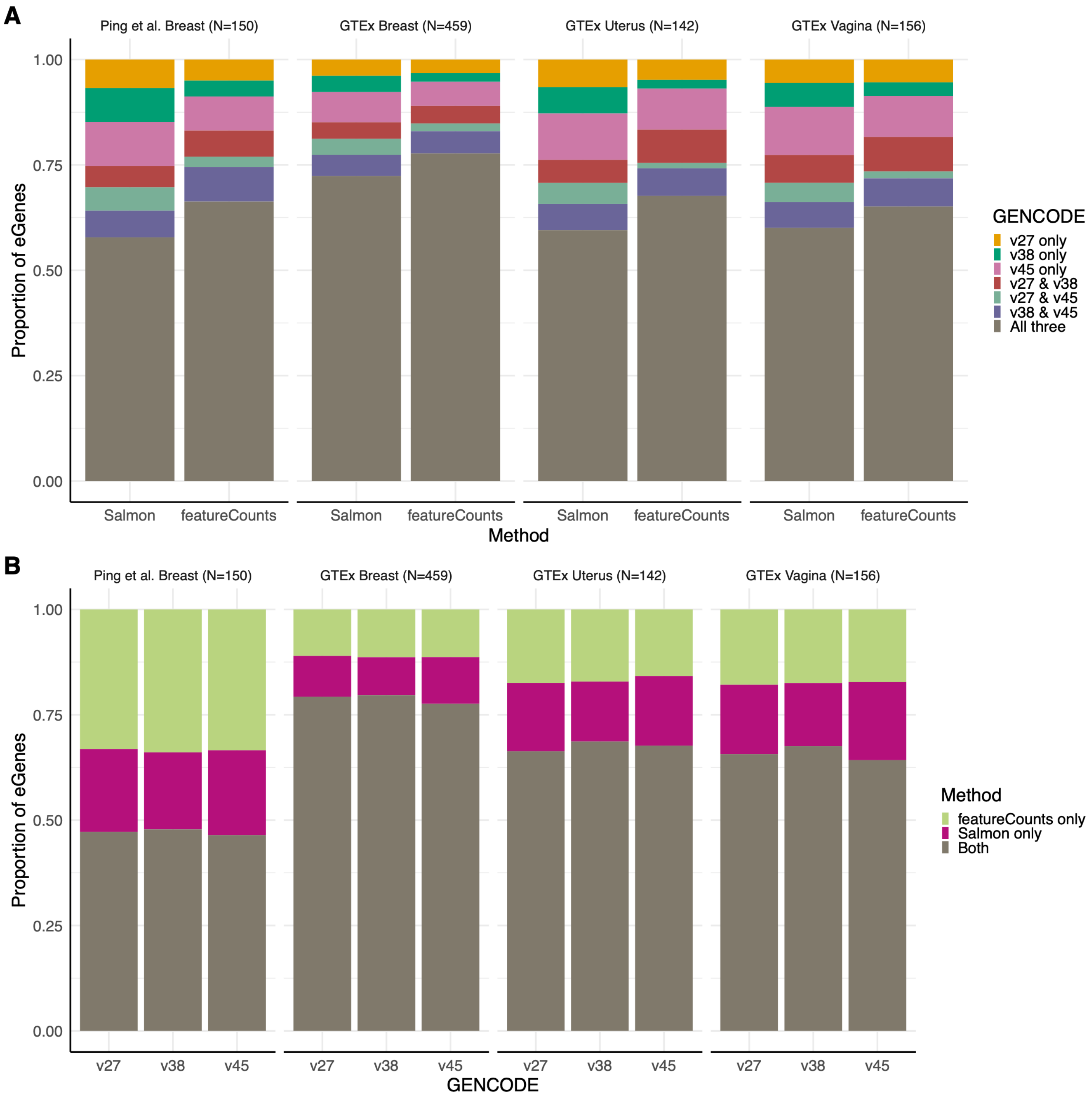


**Figure S3. (A)** Proportion of eGenes identified by one or more of the three GENCODE annotation versions, for a given method, in breast tissue samples from Ping et al. (left), breast tissue samples from GTEx (middle left), and two other non-brain tissue types from GTEx most similar in sample size to Ping et al. (right). **(B)** Proportion of eGenes identified by Salmon, STAR with featureCounts, or both, for a given annotation, across these four sample sets.

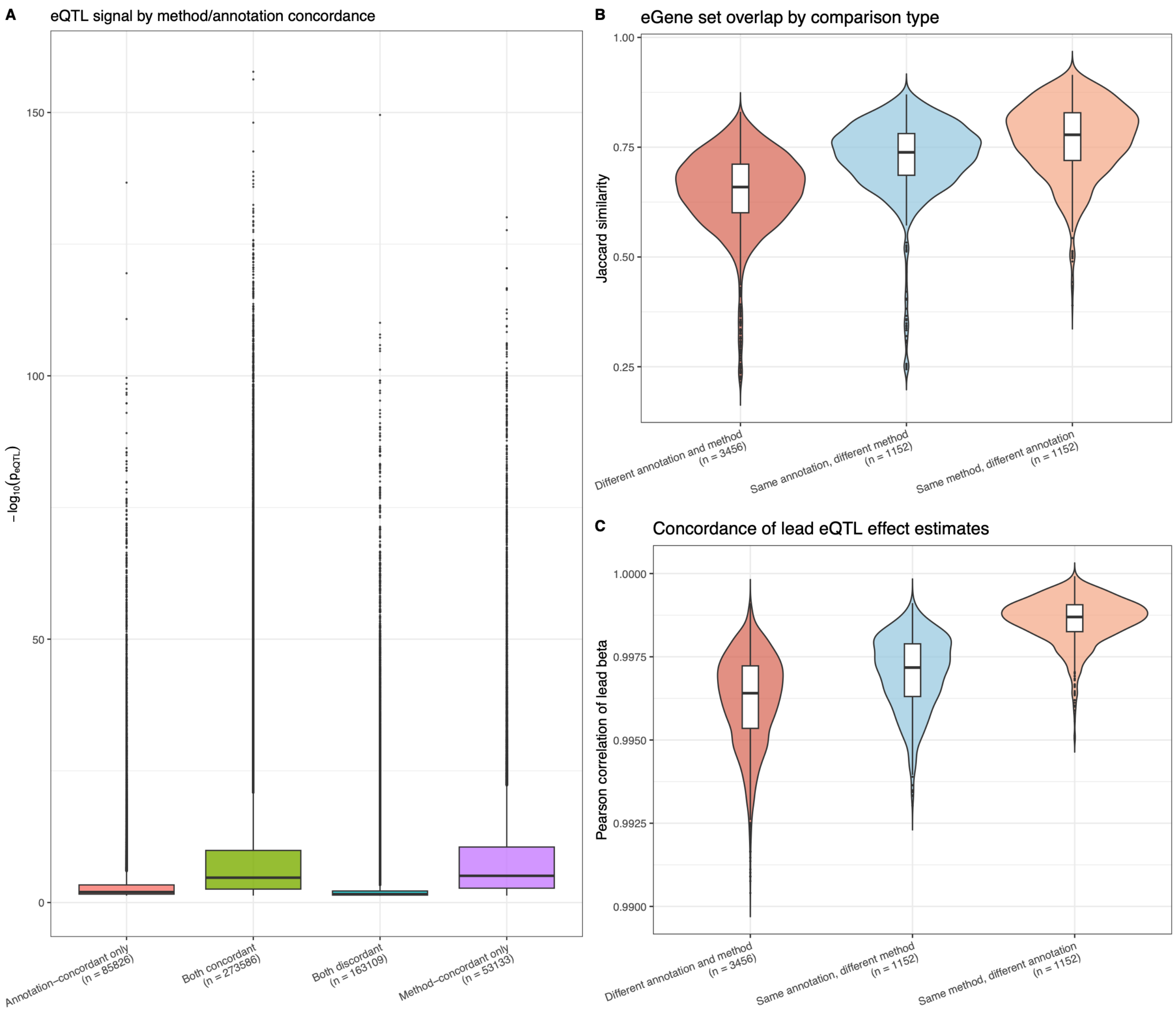


**Figure S4.** QTL signal strength and effect size concordance across quantification method and annotation comparisons. (**A**) Distribution of mean $-\log_{10}$(lead eQTL p-value) across eGenes stratified by concordance type (both concordant, method-concordant only, annotation-concordant only, both discordant). (**B**) Pairwise Jaccard similarity of eGene sets and (**C**) Pearson correlation of lead eQTL effect sizes for shared lead eQTLs across runs with different annotation/method, shared annotation, and shared method, pooled across 48 GTEx tissues. The whiskers extend to the most extreme points within 1.5*IQR. Points beyond them are plotted individually.

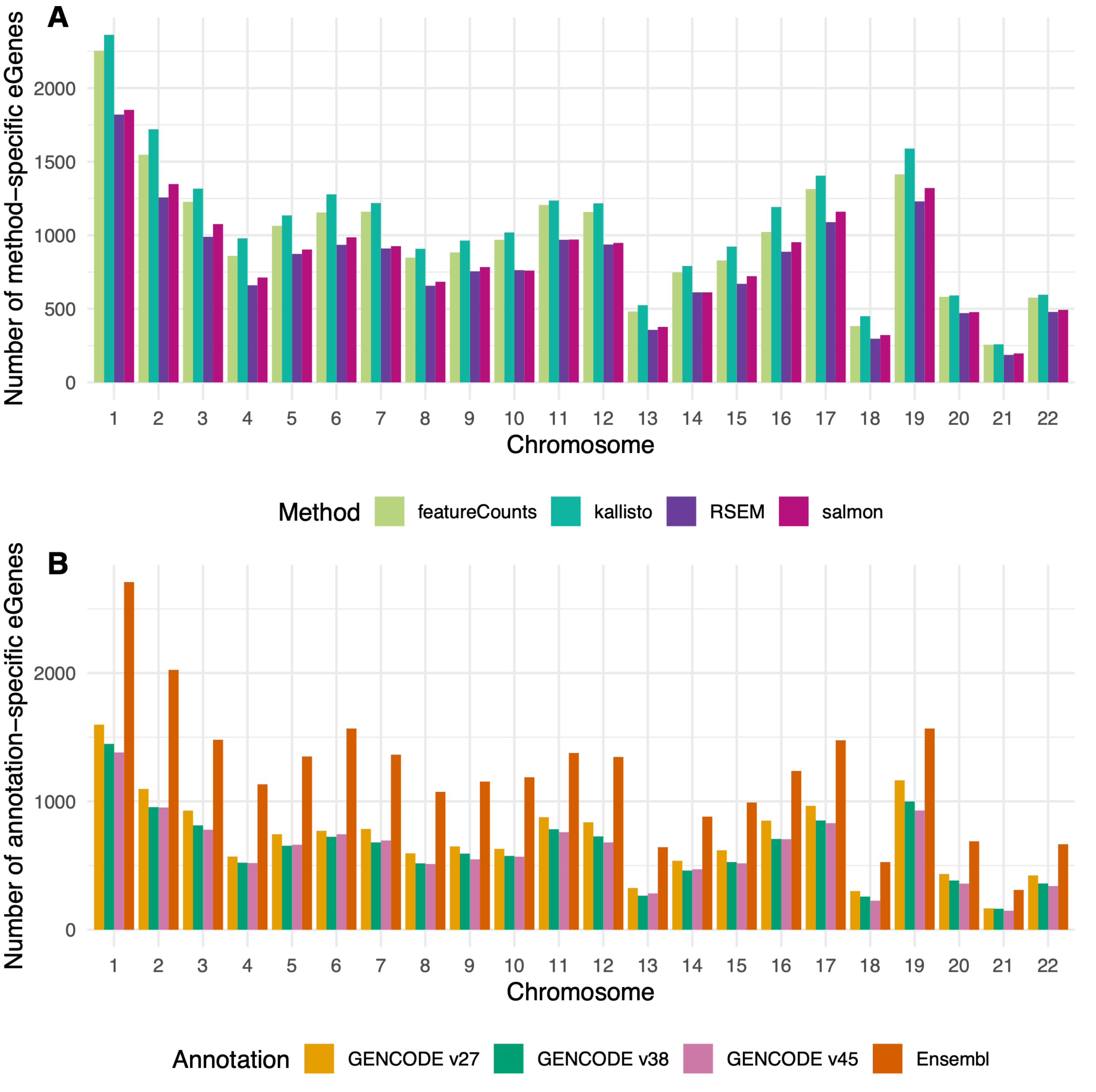


**Figure S5:** Number of eGenes by chromosome uniquely identified in a dataset specific to **(A)** quantification method and **(B)** transcriptome annotation.

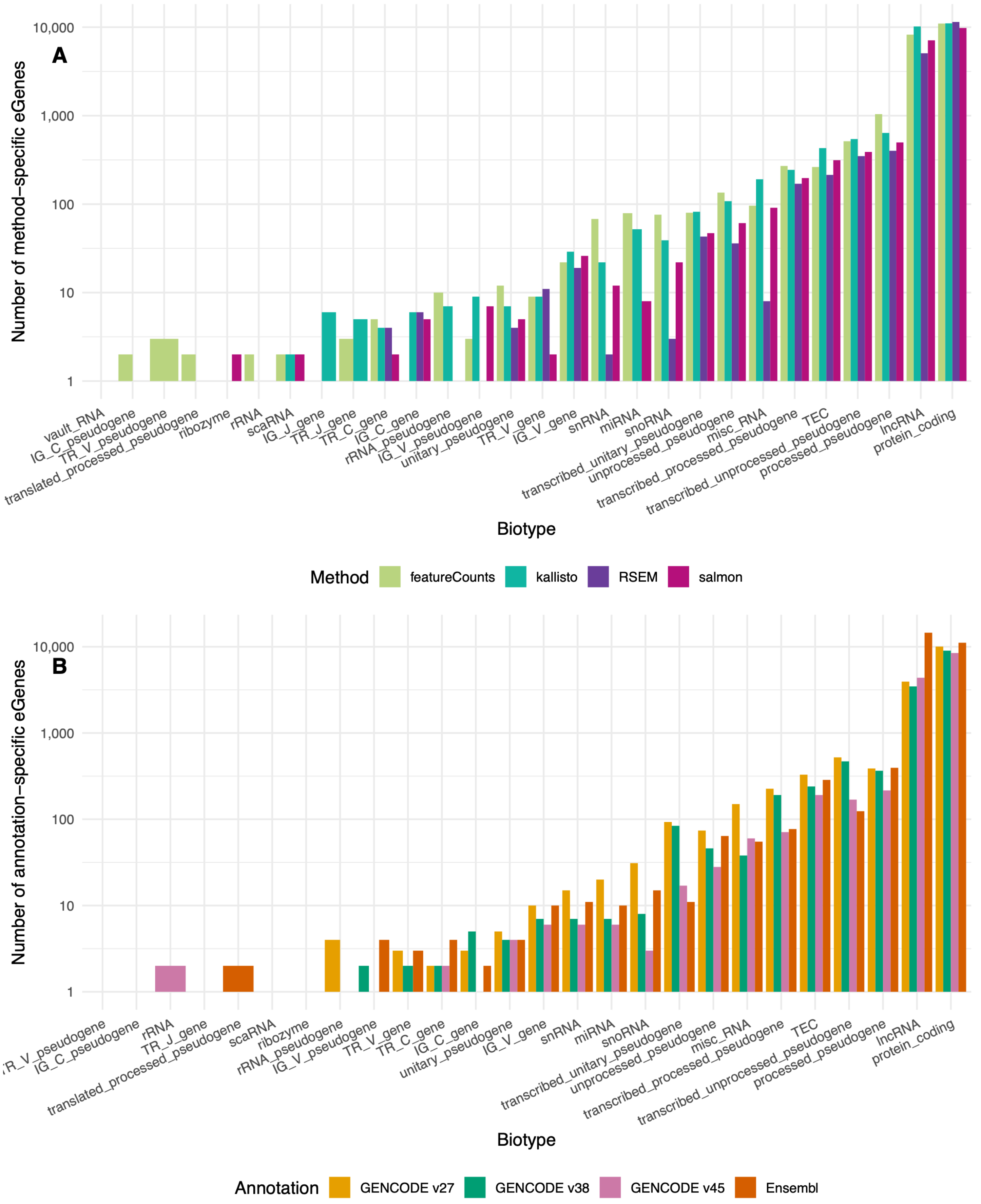


**Figure S6:** Number of eGenes by biotype uniquely identified in a dataset specific to **(A)** quantification method and **(B)** transcriptome annotation.

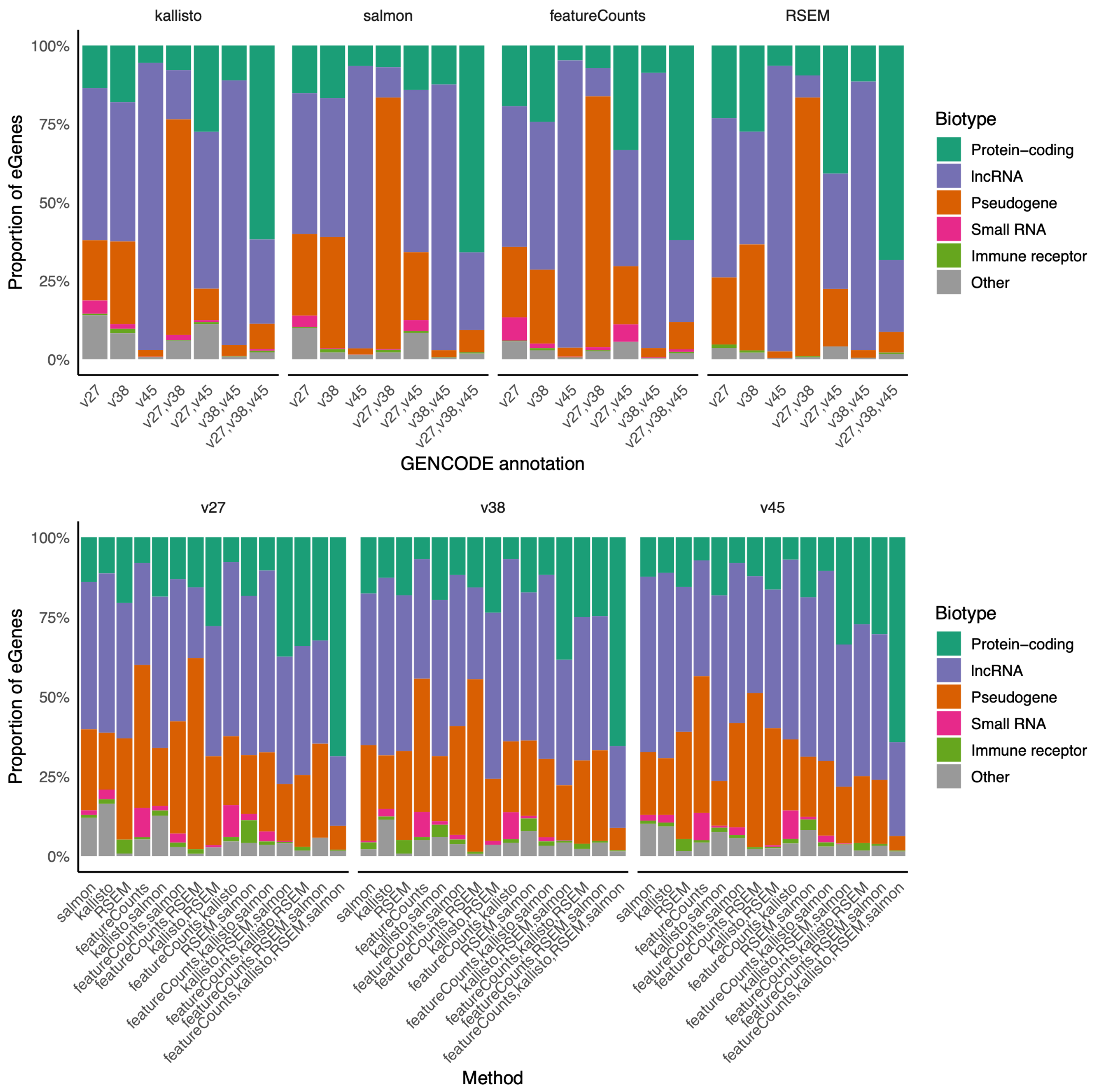


**Figure S7: Top:** Distribution of biotype in eGenes identified by different combinations of annotations (x-axis, here, exclusively considered GENCODE versions) in a dataset specific to a quantification method (vertical panels). The furthest right bar within each panel represents those eGenes identified by all three GENCODE annotations, while the first three bars represent those eGenes identified in only one. **Bottom:** Distribution of biotype in eGenes identified by different combinations of quantification methods (x-axis) in a dataset specific to a GENCODE annotation (vertical panels). The furthest right bar within each panel represents those eGenes identified by all four methods, while the first four bars represent those eGenes identified in only one.

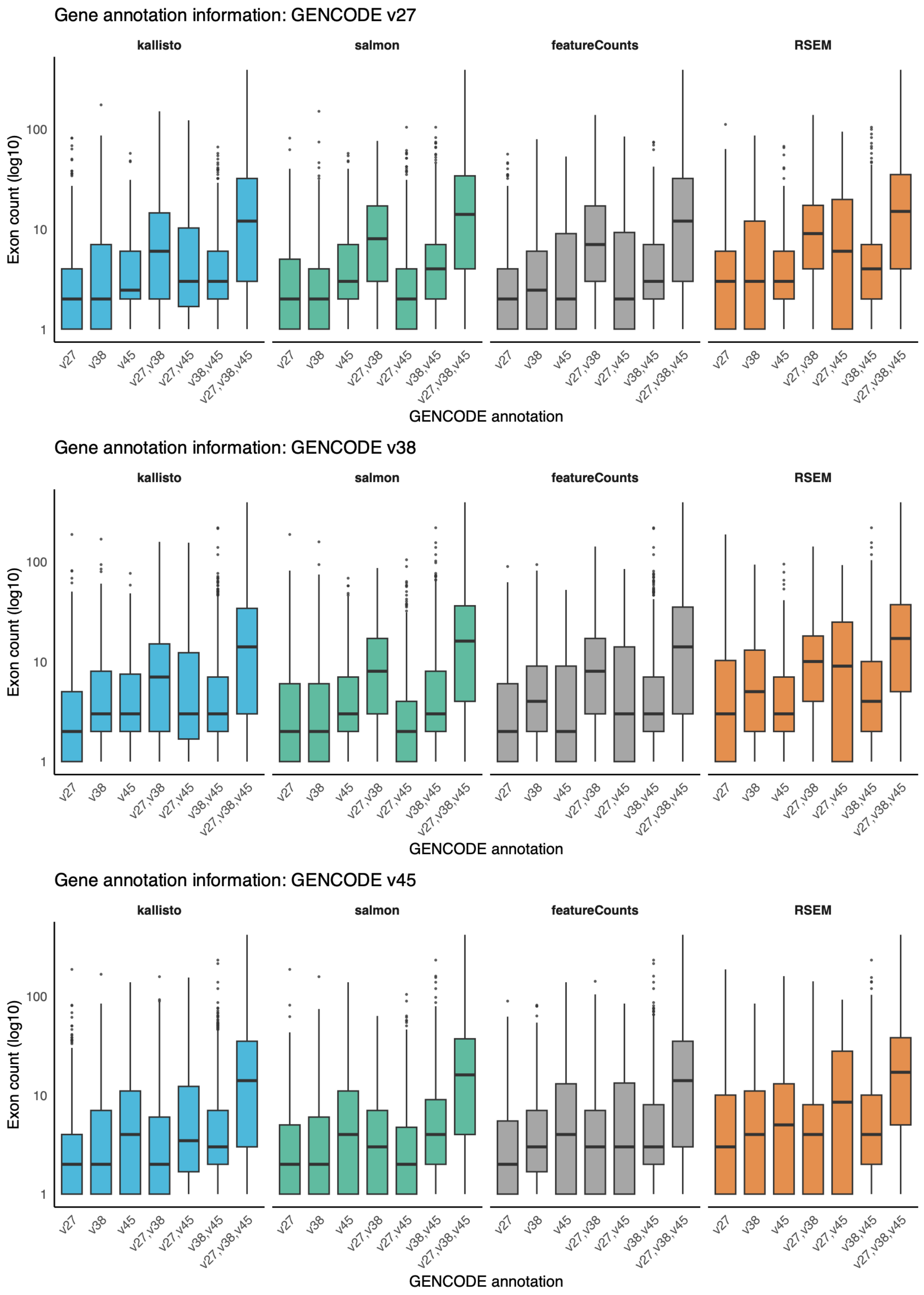


**Figure S8:** Distribution of number of exons in eGenes identified by different combinations of annotations (x-axis, here, exclusively considered GENCODE versions) in a dataset specific to a quantification method (vertical panels). The furthest right boxplot within each panel represents those eGenes identified by all three GENCODE annotations, while the first three boxplots represent those eGenes identified in only one. The horizontal panels indicate which GENCODE annotation was used to calculate the number of exons per gene. The whiskers extend to the most extreme points within 1.5*IQR. Points beyond them are plotted individually.

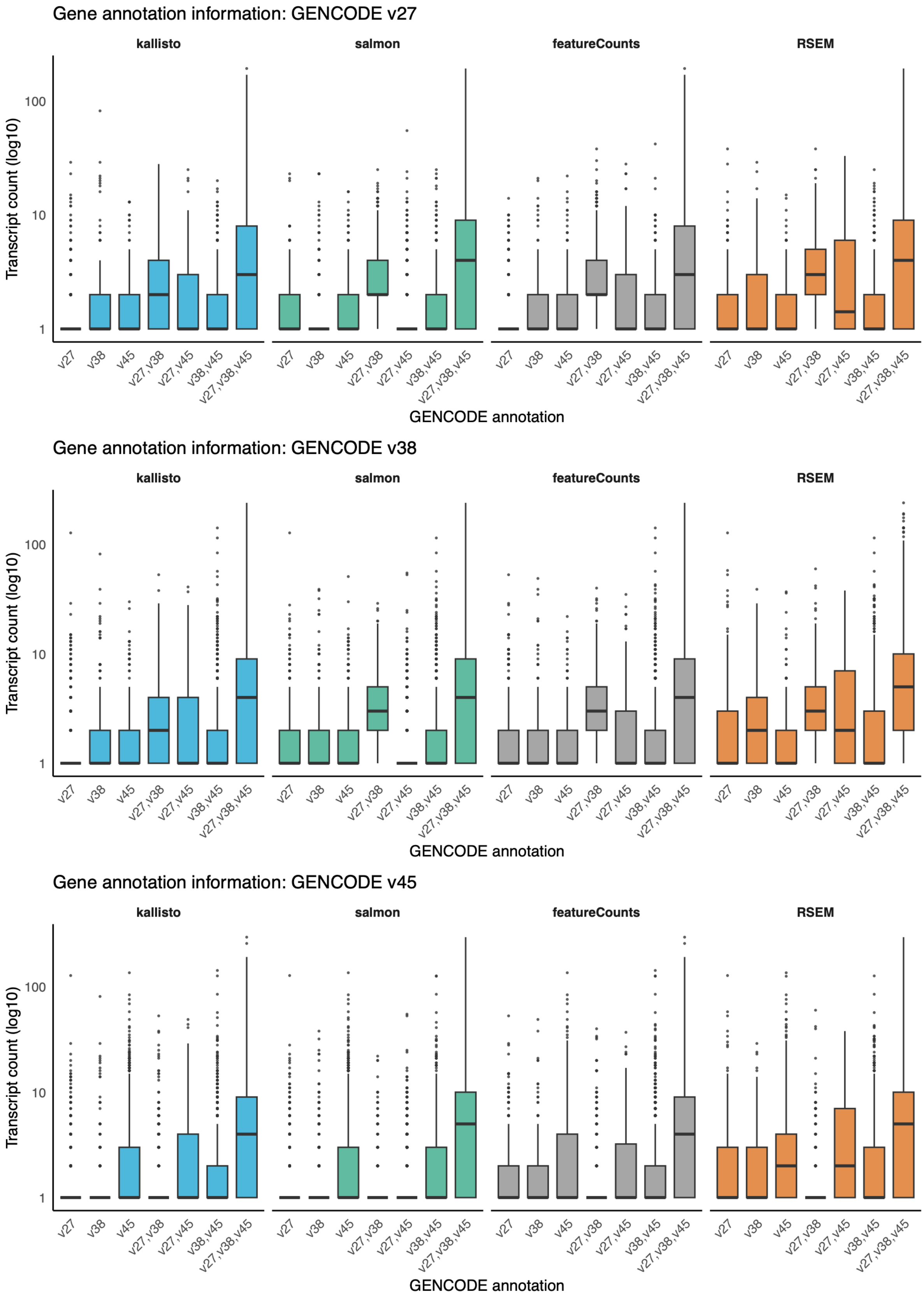


**Figure S9:** Distribution of number of transcripts in eGenes identified by different combinations of GENCODE annotations (x-axis) in a dataset specific to a quantification method (vertical panels). The furthest right boxplot within each panel represents those eGenes identified by all three annotations, while the first three boxplots represent those eGenes identified in only one. The horizontal panels indicate which GENCODE annotation was used to calculate the number of transcripts per gene. The whiskers extend to the most extreme points within 1.5*IQR. Points beyond them are plotted individually.

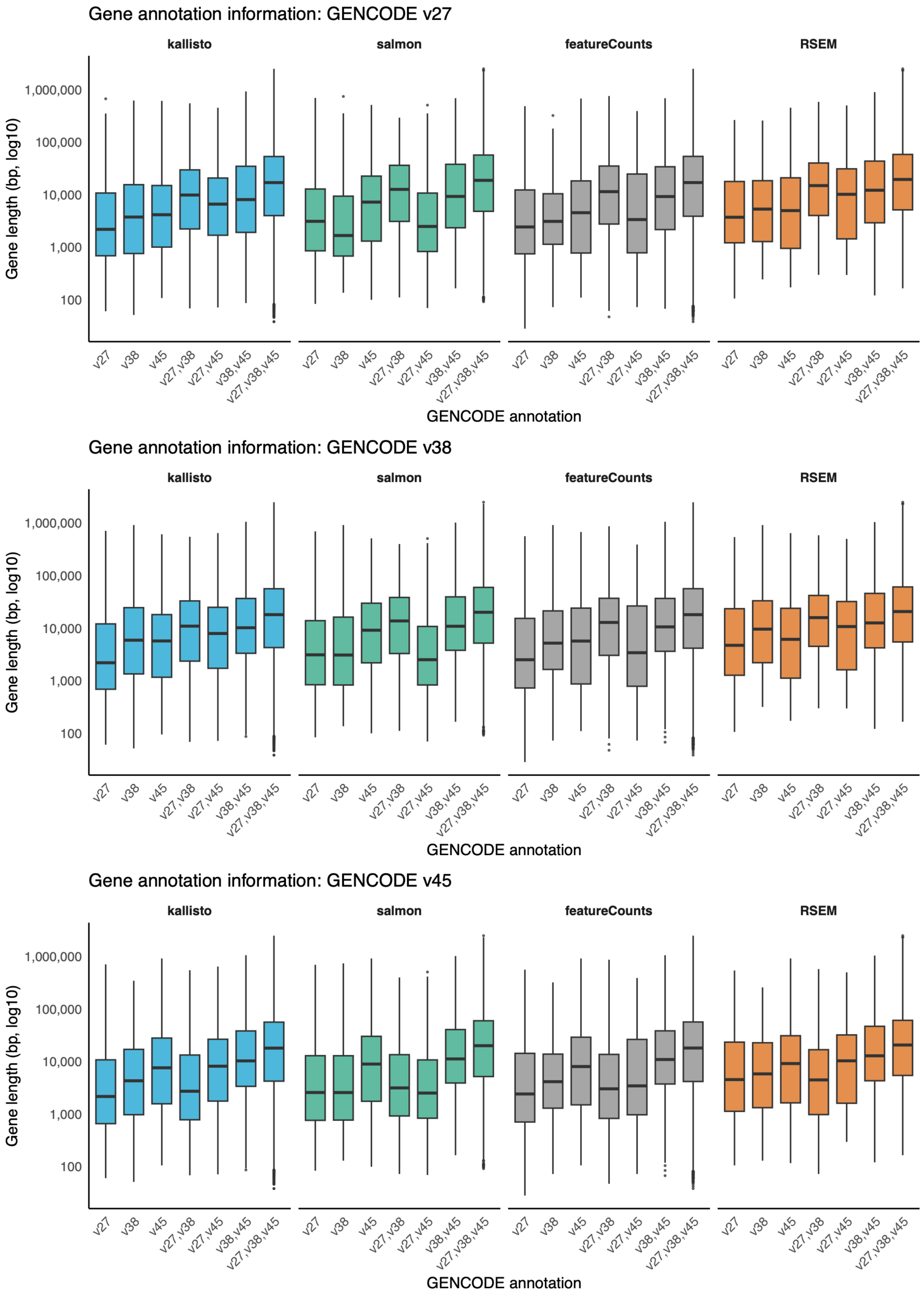


**Figure S10:** Distribution of gene length for eGenes identified by different combinations of GENCODE annotations (x-axis) in a dataset specific to a quantification method (vertical panels). The furthest right boxplot within each panel represents those eGenes identified by all three annotations, while the first three boxplots represent those eGenes identified in only one. The horizontal panels indicate which GENCODE annotation was used to calculate the gene length. The whiskers extend to the most extreme points within 1.5*IQR. Points beyond them are plotted individually.

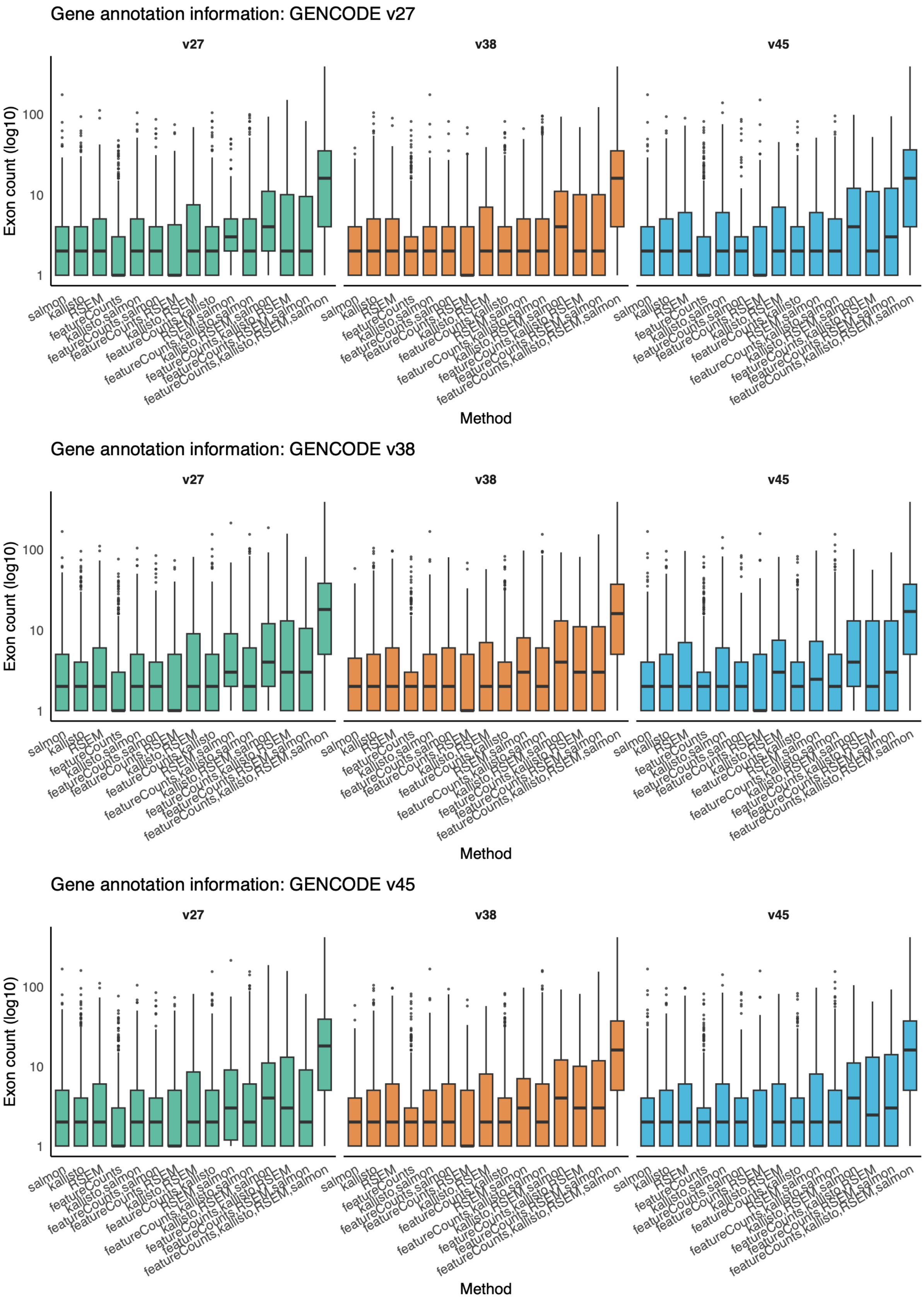


**Figure S11:** Distribution of number of exons in eGenes identified by different combinations of quantification methods (x-axis) in a dataset specific to a GENCODE transcript annotation (vertical panels). The furthest right boxplot within each panel represents those eGenes identified by all four quantification methods, while the first four boxplots represent those eGenes identified in only one. The horizontal panels indicate which GENCODE annotation was used to calculate the number of exons per gene. The whiskers extend to the most extreme points within 1.5*IQR. Points beyond them are plotted individually.

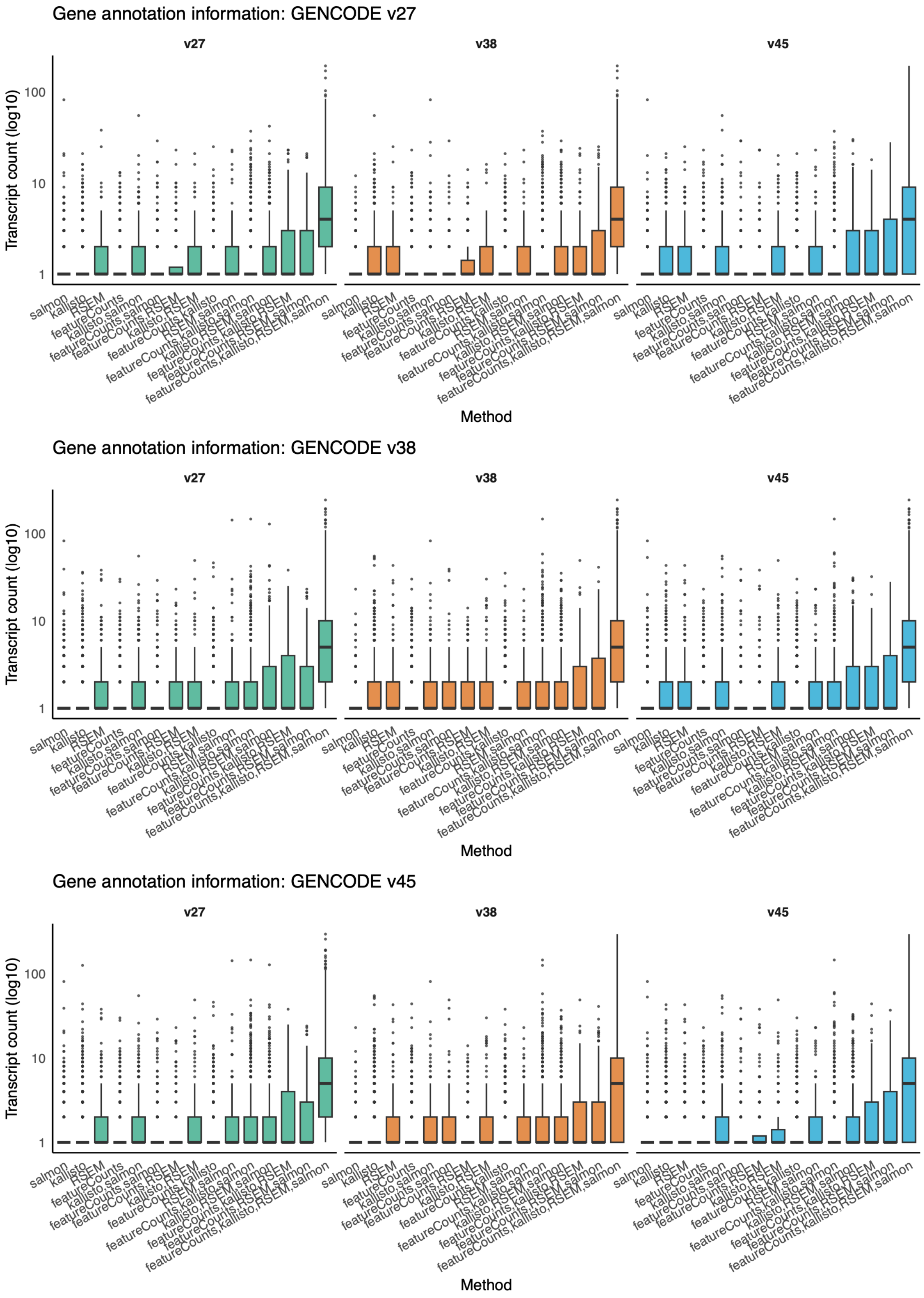


**Figure S12:** Distribution of number of transcripts in eGenes identified by different combinations of quantification methods (x-axis) in a dataset specific to a GENCODE transcript annotation (vertical panels). The furthest right boxplot within each panel represents those eGenes identified by all four quantification methods, while the first four boxplots represent those eGenes identified in only one. The horizontal panels indicate which GENCODE annotation was used to calculate the number of transcripts per gene. The whiskers extend to the most extreme points within 1.5*IQR. Points beyond them are plotted individually.

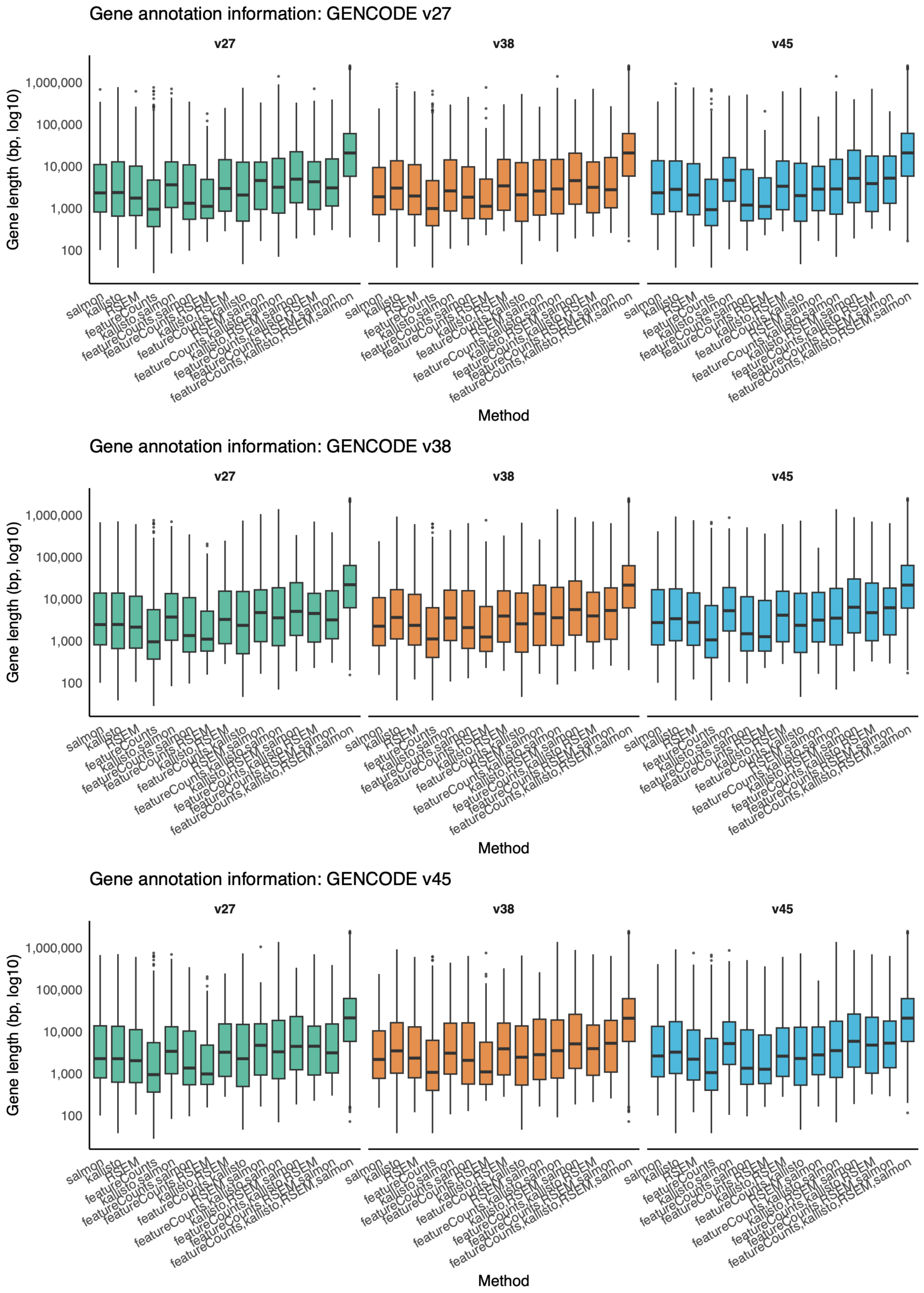


**Figure S13:** Distribution of number of transcripts in eGenes identified by different combinations of quantification methods (x-axis) in a dataset specific to a GENCODE transcript annotation (vertical panels). The furthest right boxplot within each panel represents those eGenes identified by all four quantification methods, while the first four boxplots represent those eGenes identified in only one. The horizontal panels indicate which GENCODE annotation was used to calculate the number of transcripts. The whiskers extend to the most extreme points within 1.5*IQR. Points beyond them are plotted individually.

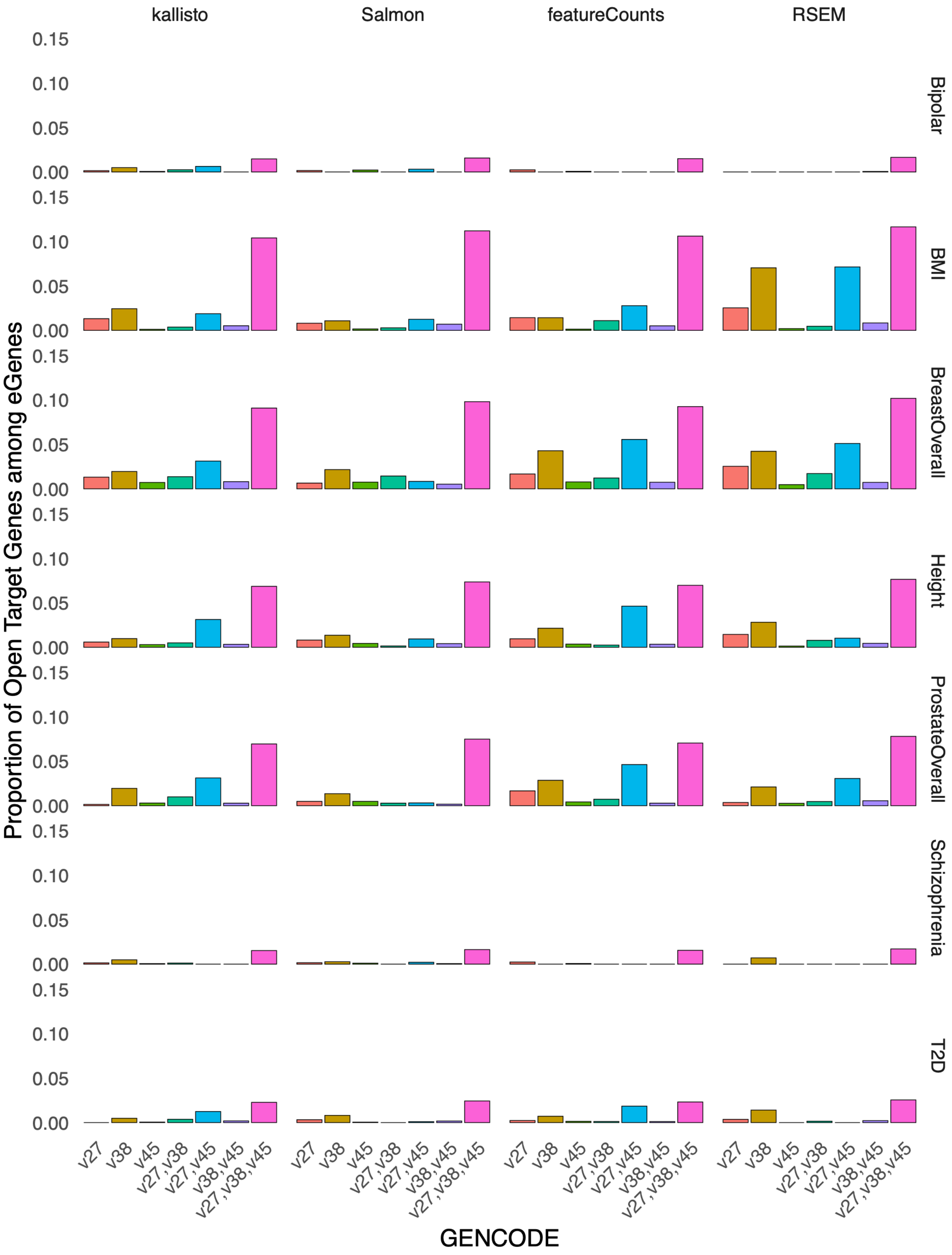


**Figure S14:** Proportion of Open Targets genes among eGenes identified by different combinations of GENCODE annotations (x-axis) in a dataset specific to a quantification method (vertical panels). The furthest right bar within each panel represents those eGenes identified by all three annotations, while the first three bars represent those eGenes identified in only one. The horizontal panels indicate which phenotype for which Open Targets was queried.

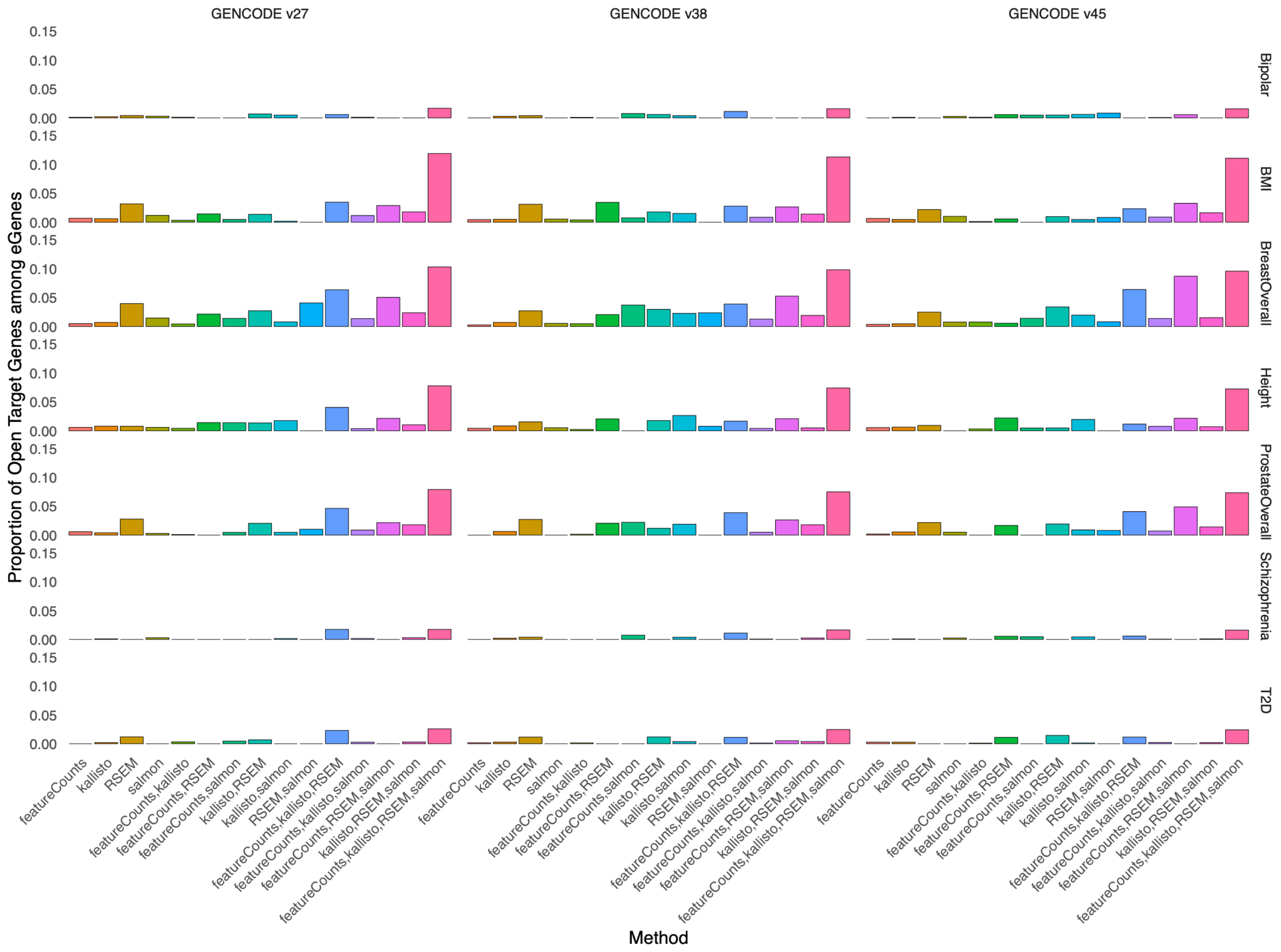


**Figure S15:** Proportion of Open Targets genes among eGenes identified by different combinations of quantification methods (x-axis) in a dataset specific to a GENCODE annotation version (vertical panels). The furthest right bar within each panel represents those eGenes identified by all four methods, while the first four bars represent those eGenes identified in only one. The horizontal panels indicate which phenotype for which Open Targets was queried.

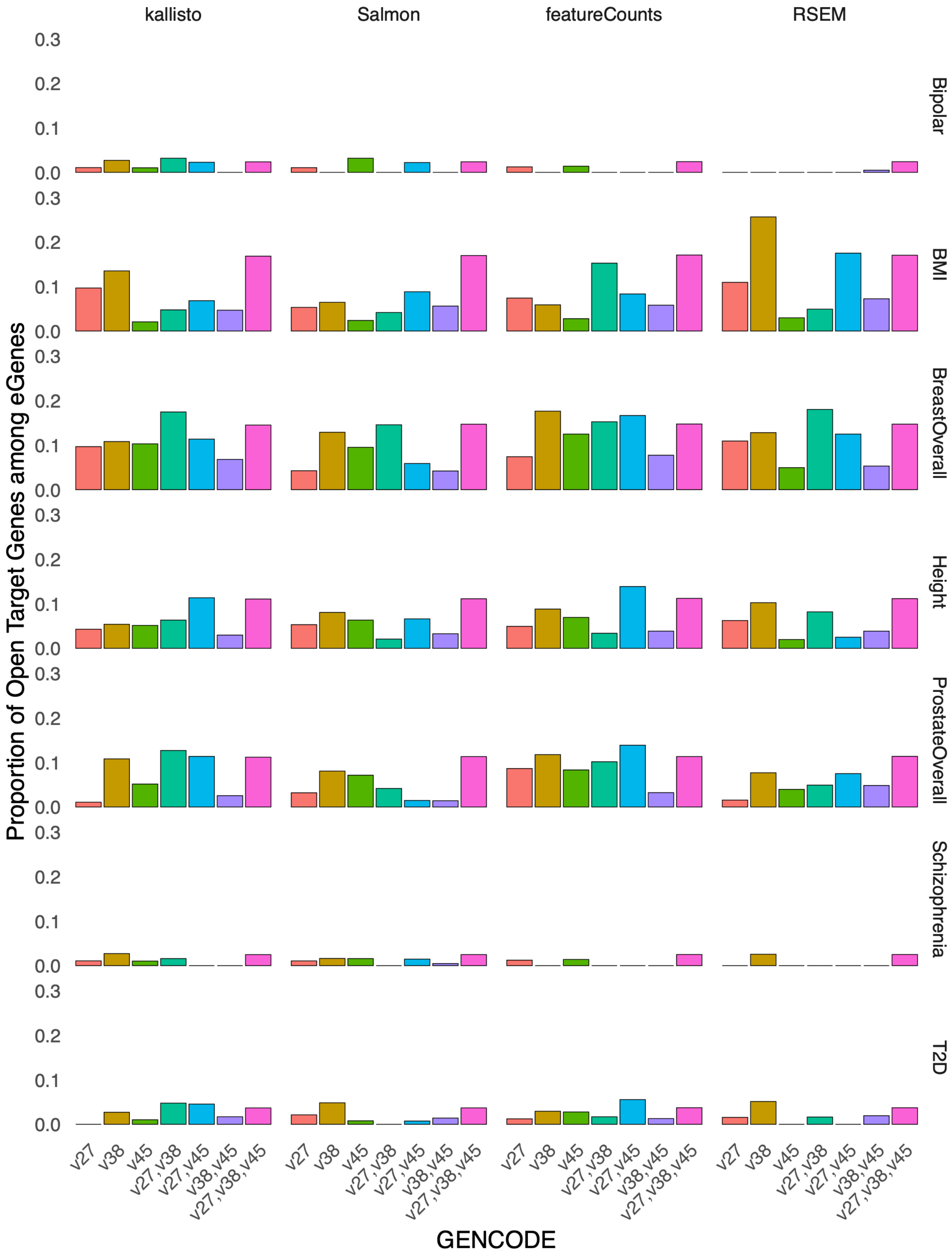


**Figure S16:** Proportion of Open Targets genes among eGenes (restricted to protein coding only) identified by different combinations of GENCODE annotations (x-axis) in a dataset specific to a quantification method (vertical panels). The furthest right bar within each panel represents those eGenes identified by all three annotations, while the first three bars represent those eGenes identified in only one. The horizontal panels indicate which phenotype for which Open Targets was queried.

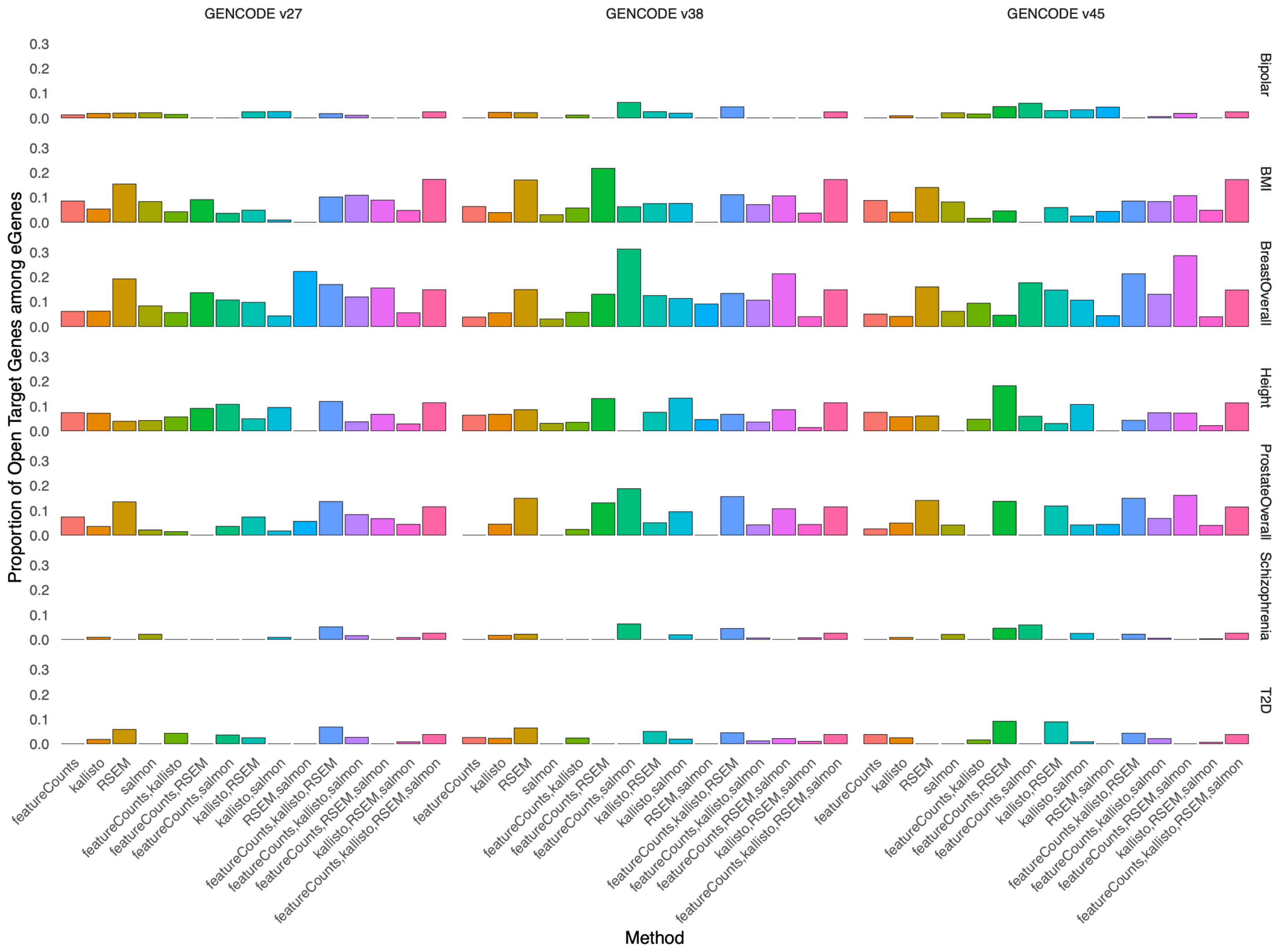


**Figure S17:** Proportion of Open Targets genes among eGenes (restricted to protein-coding only) identified by different combinations of quantification methods (x-axis) in a dataset specific to a GENCODE annotation version (vertical panels). The furthest right bar within each panel represents those eGenes identified by all four methods, while the first four bars represent those eGenes identified in only one. The horizontal panels indicate which phenotype for which Open Targets was queried.

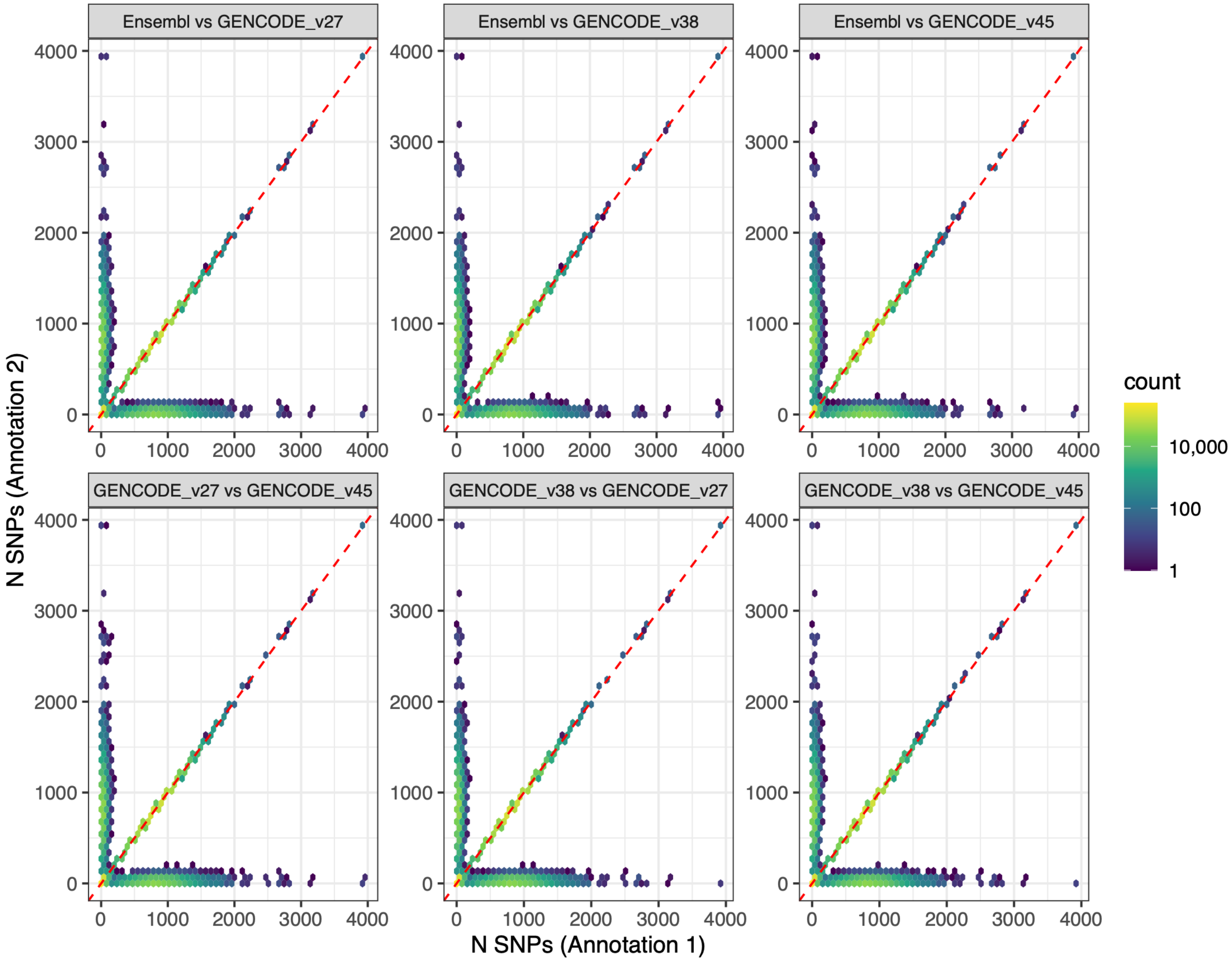


**Figure S18:** Hexagonal density plots of number of SNPs in expression prediction models for the same gene in the same tissue with the same quantification method across transcriptome annotations.

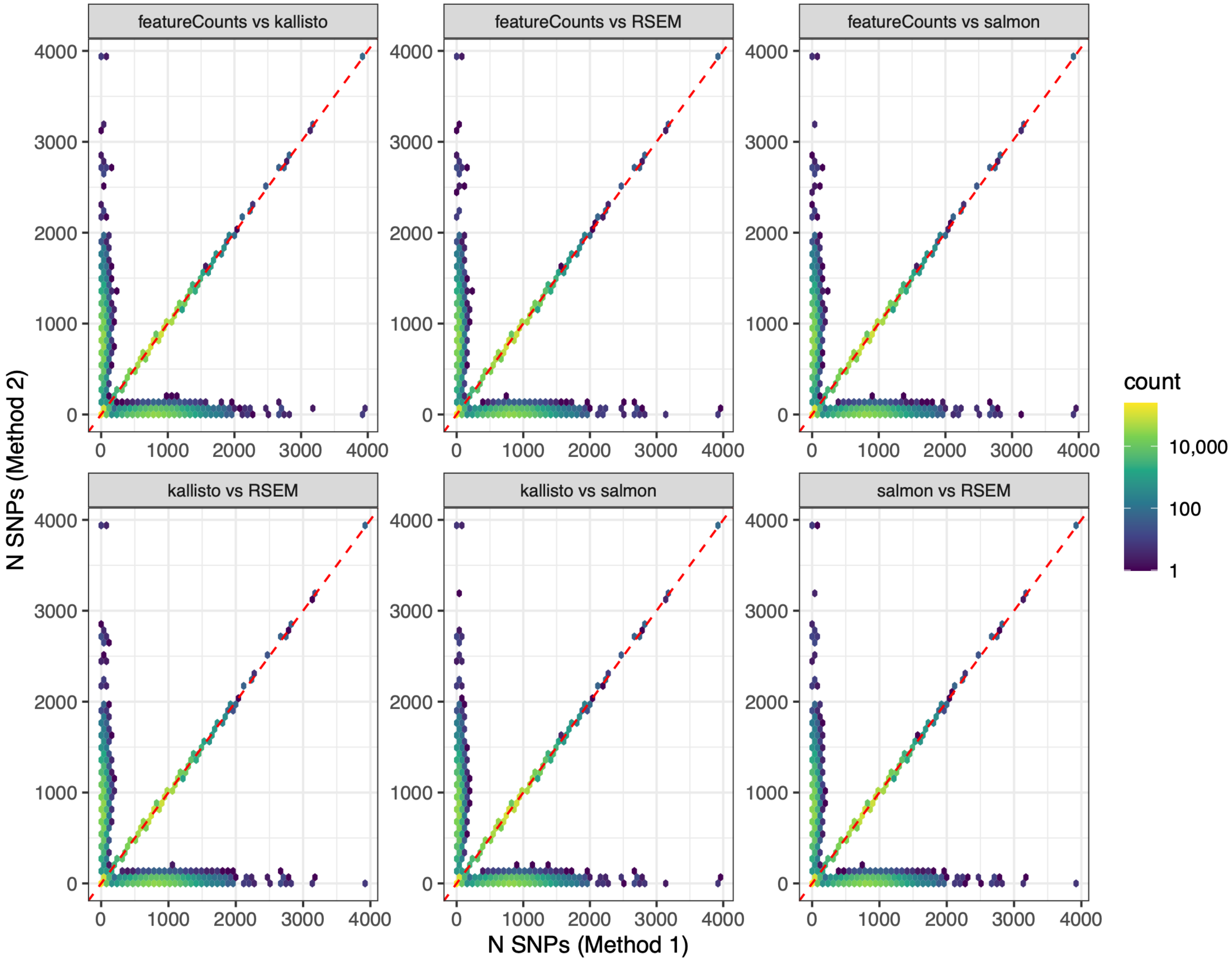


**Figure S19:** Hexagonal density plots of number of SNPs in expression prediction models for the same gene in the same tissue with the same annotation across quantification methods.

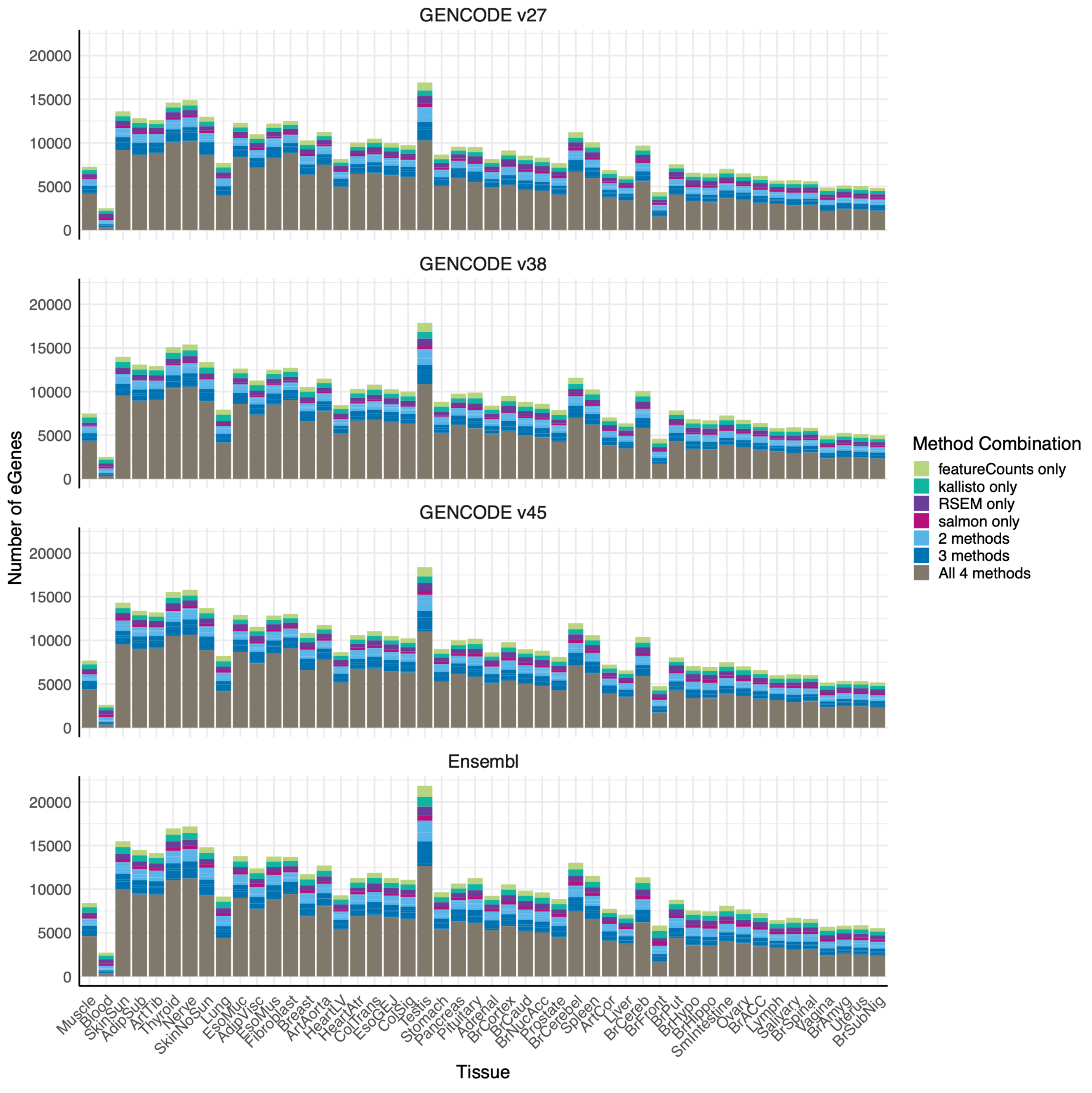


**Figure S20**: Number of eGenes (Y-axis) called at adjusted P < 0.05 by method across each GTEx tissue and annotation, arranged in decreasing sample size order (X-axis).

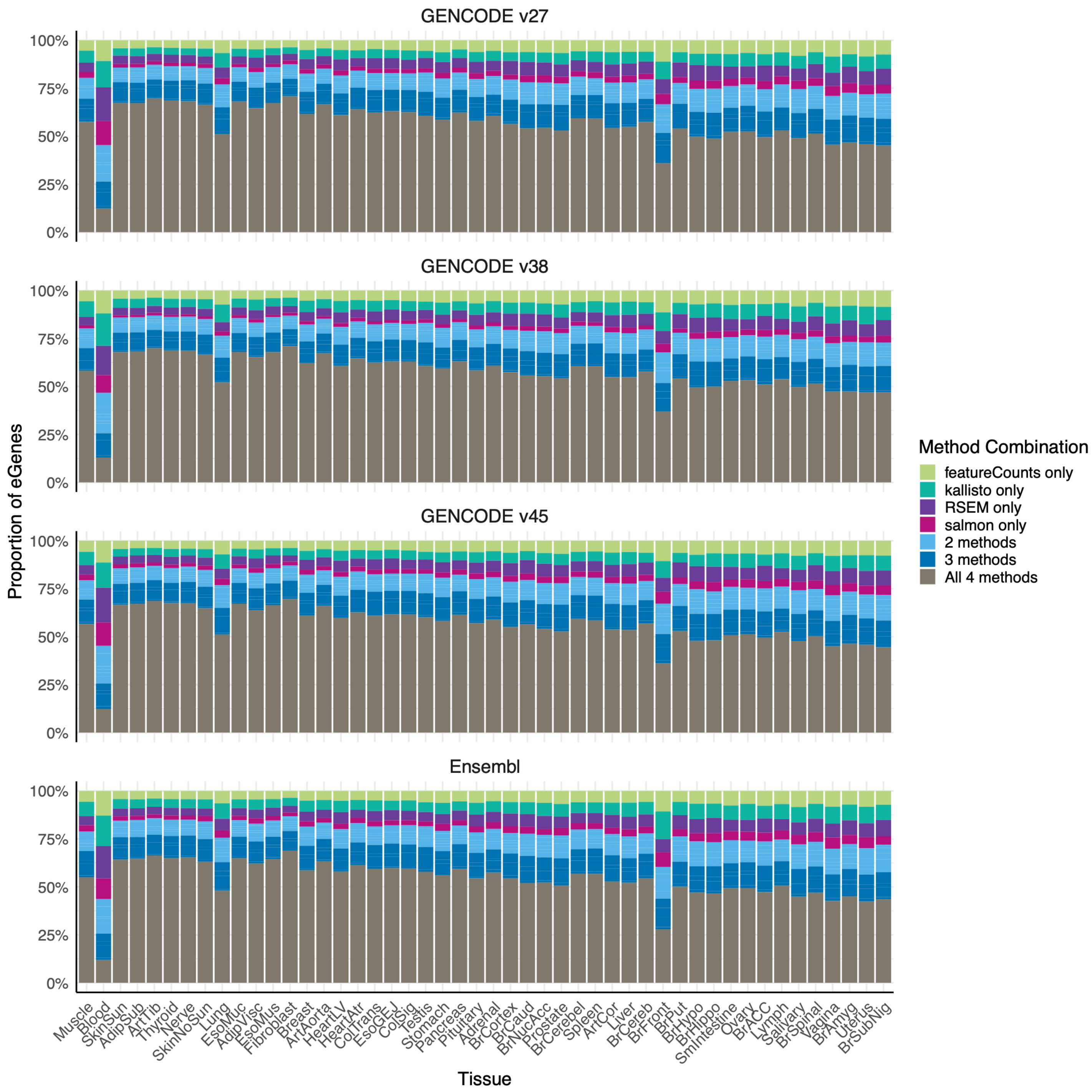


**Figure S21**: Proportion of eGenes (Y-axis) called at adjusted P < 0.05 by method across each GTEx tissue and annotation, arranged in decreasing sample size order (X-axis).

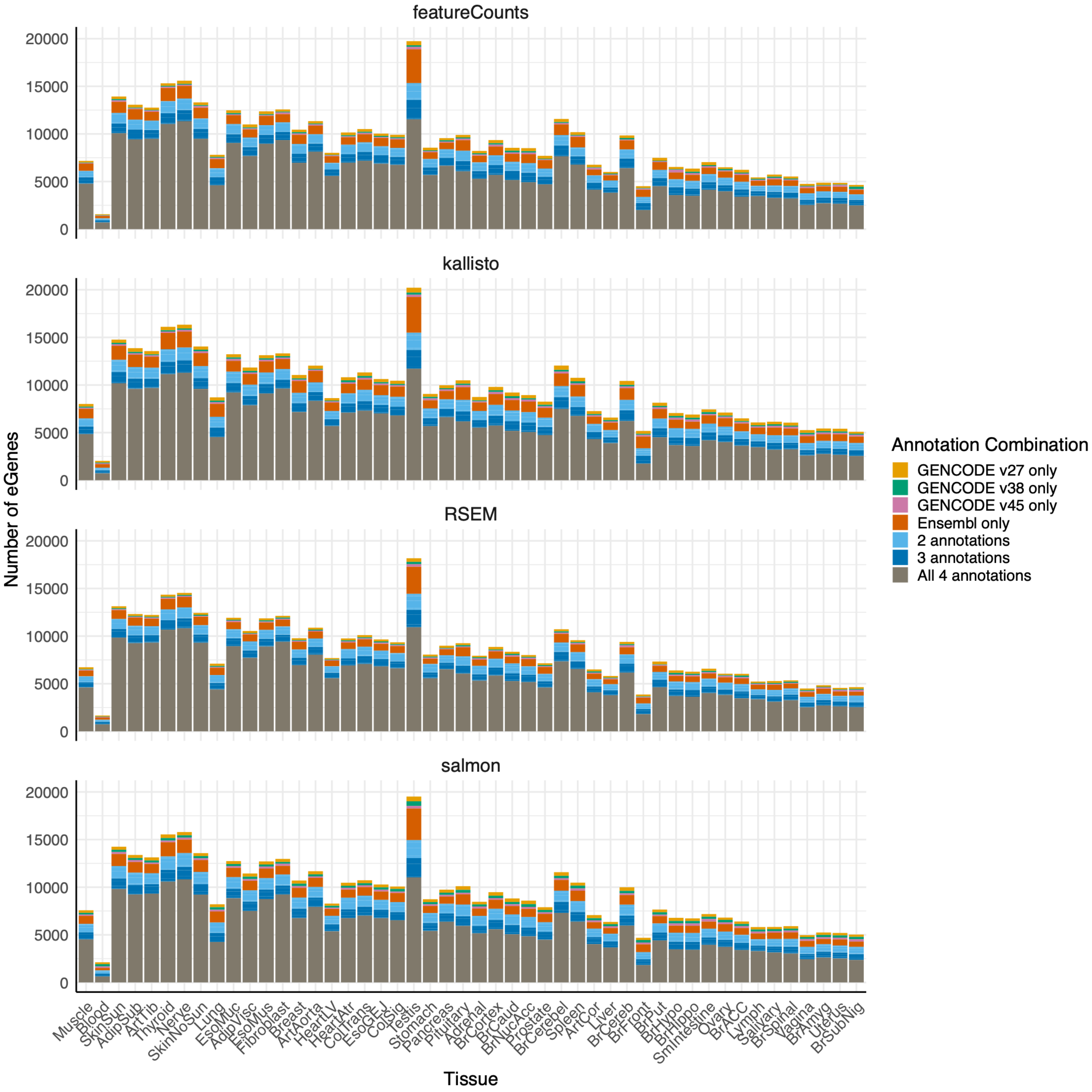


**Figure S22**: Number of eGenes (Y-axis) called at adjusted P < 0.05 by transcriptome annotation across each GTEx tissue and quantification method, arranged in decreasing sample size order (X-axis).

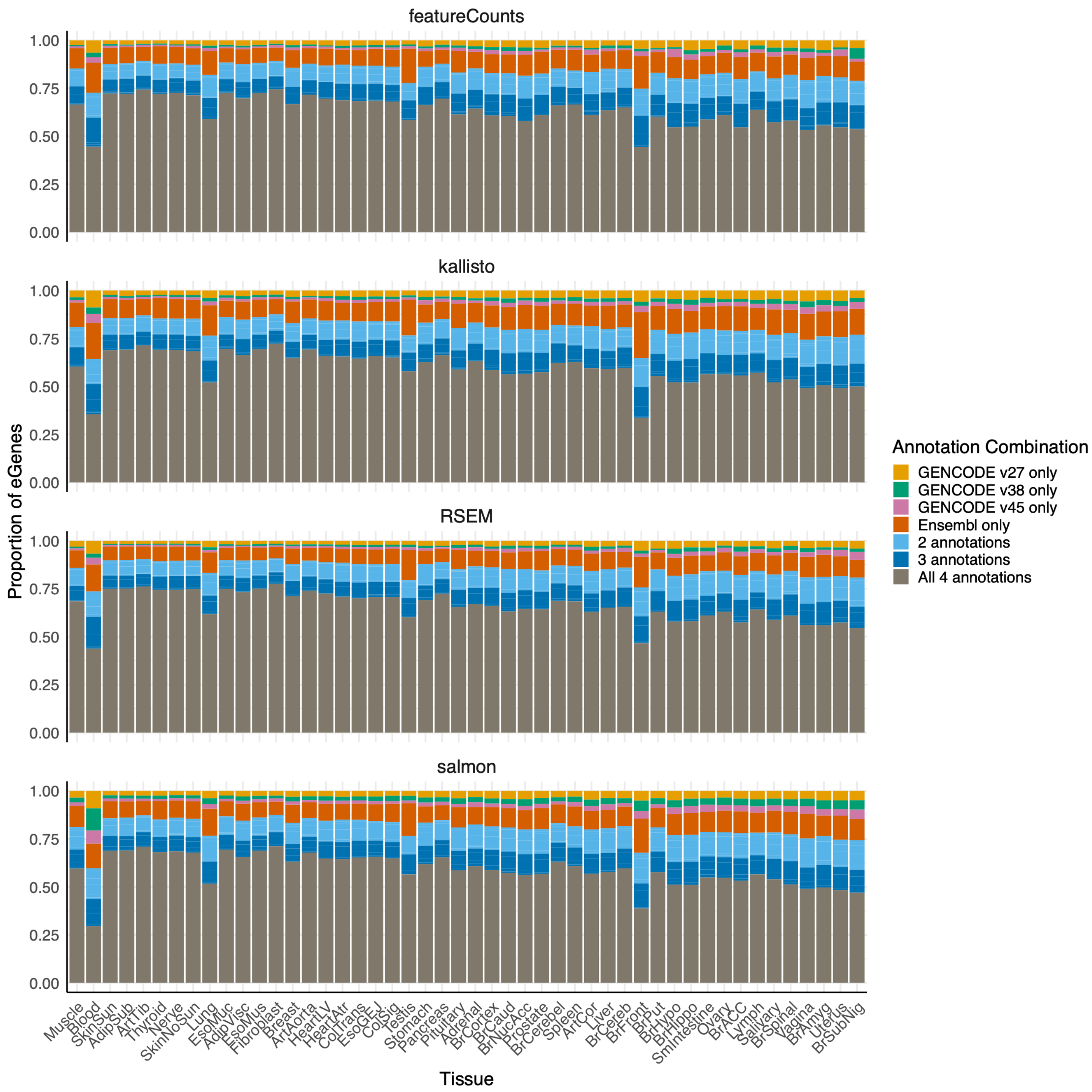


**Figure S23:** Proportion of eGenes (Y-axis) called at adjusted P < 0.05 by transcriptome annotation across each GTEx tissue and quantification method, arranged in decreasing sample size order (X-axis).

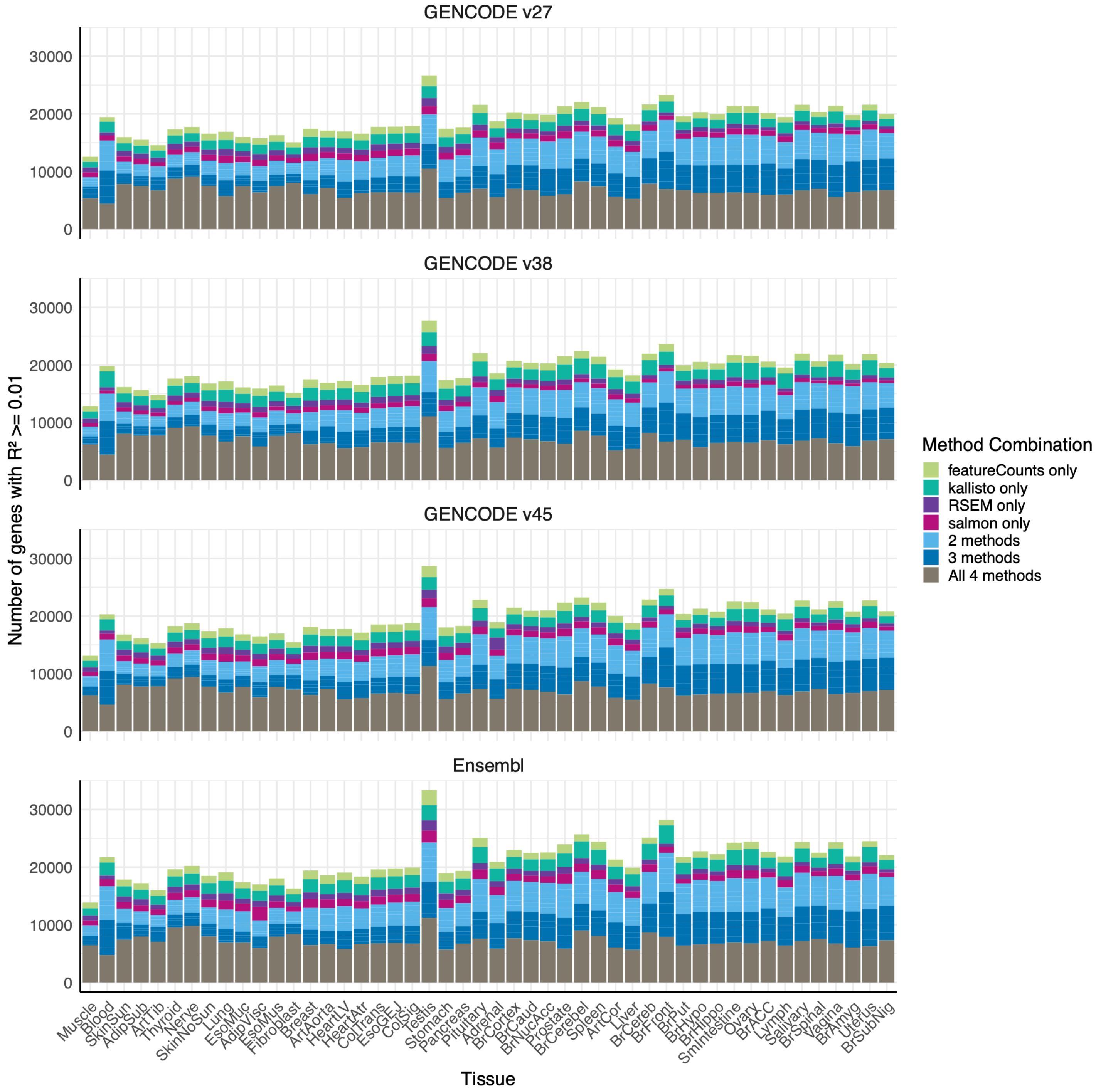


**Figure S24**: Number of genes predicted at cross-validation $R^2 \geq 0.01$ (Y-axis) called at adjusted $P < 0.05$ by method across each GTEx tissue and annotation, arranged in decreasing sample size order (X-axis).

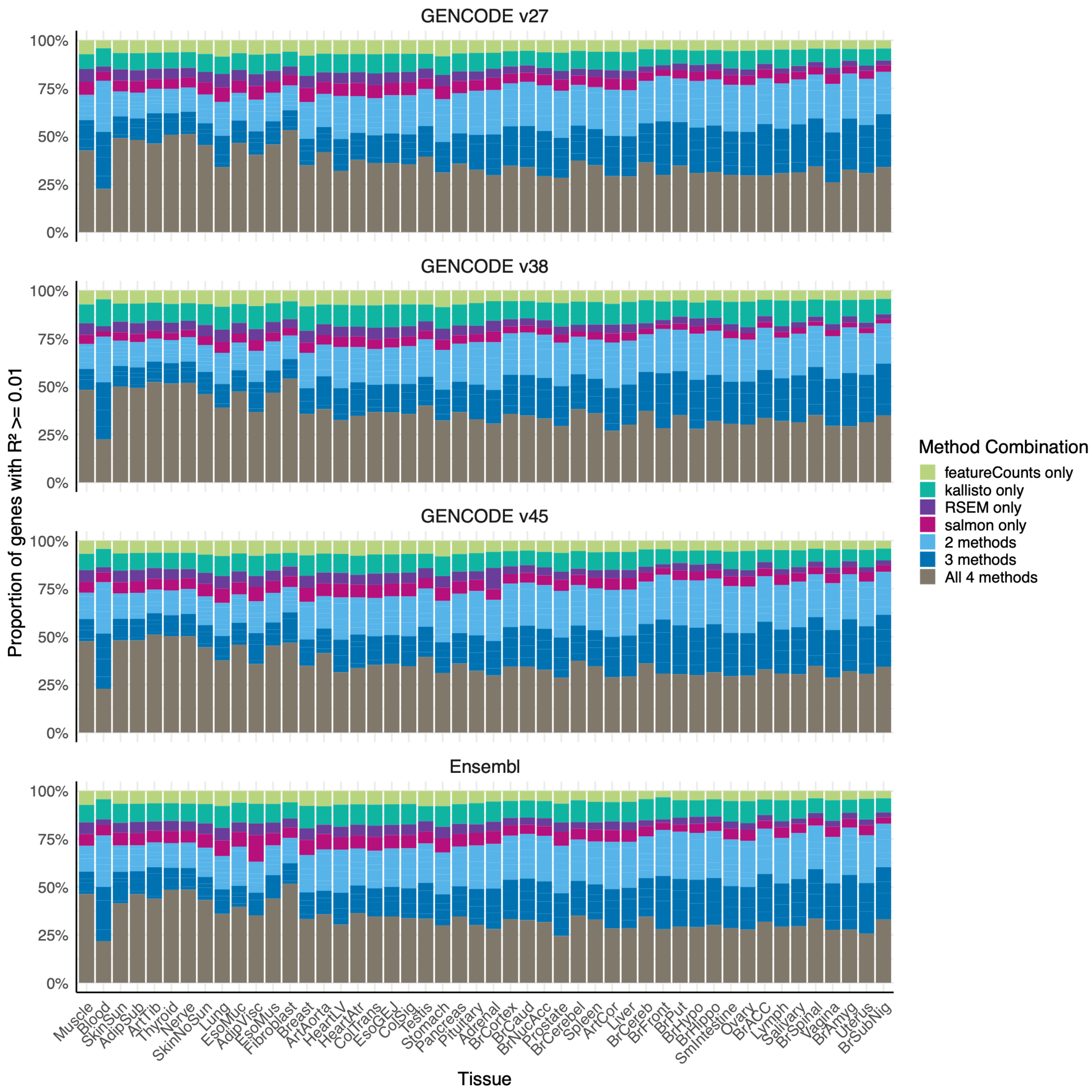


**Figure S25**: Proportion of genes predicted at cross-validation $R^2 \geq 0.01$ (Y-axis) by method across each GTEx tissue and annotation, arranged in decreasing sample size order (X-axis).

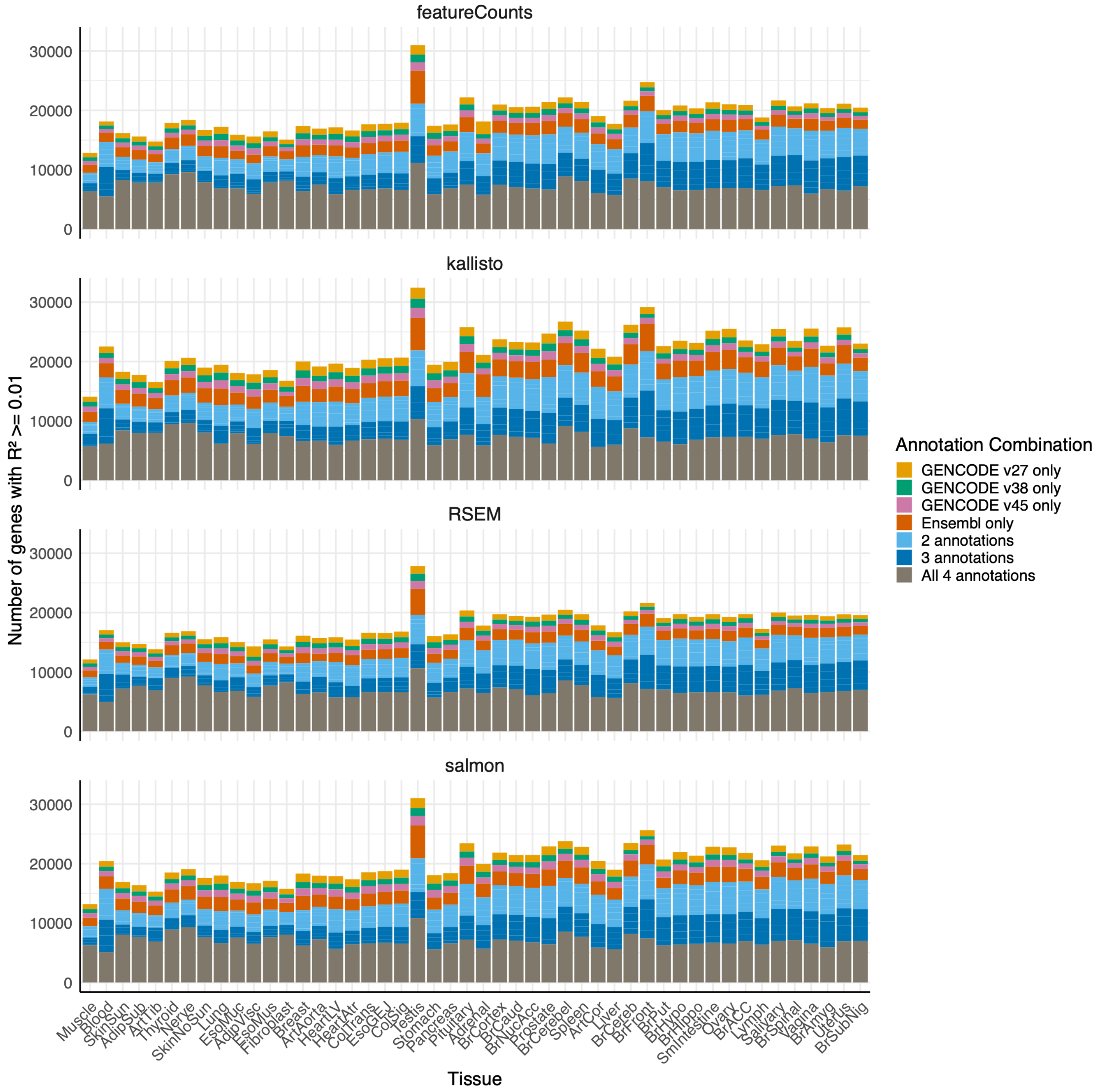


**Figure S26**: Number of genes predicted at cross-validation $R^2 \geq 0.01$ (Y-axis) by transcriptome annotation across each GTEx tissue and quantification method, arranged in decreasing sample size order (X-axis).

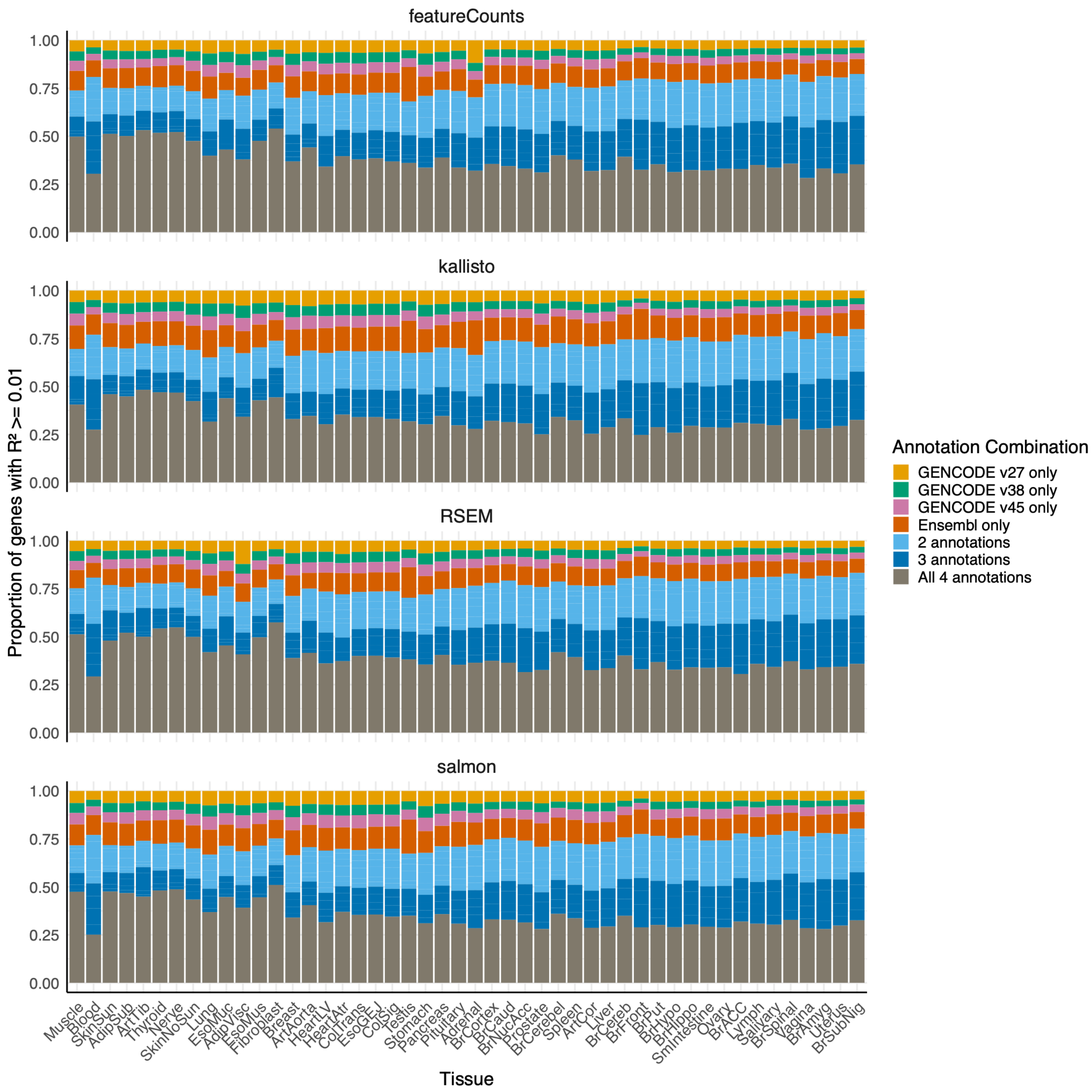


**Figure S27**: Proportion of genes predicted at cross-validation $R^2 \geq 0.01$ (Y-axis) by transcriptome annotation across each GTEx tissue and quantification method, arranged in decreasing sample size order (X-axis).

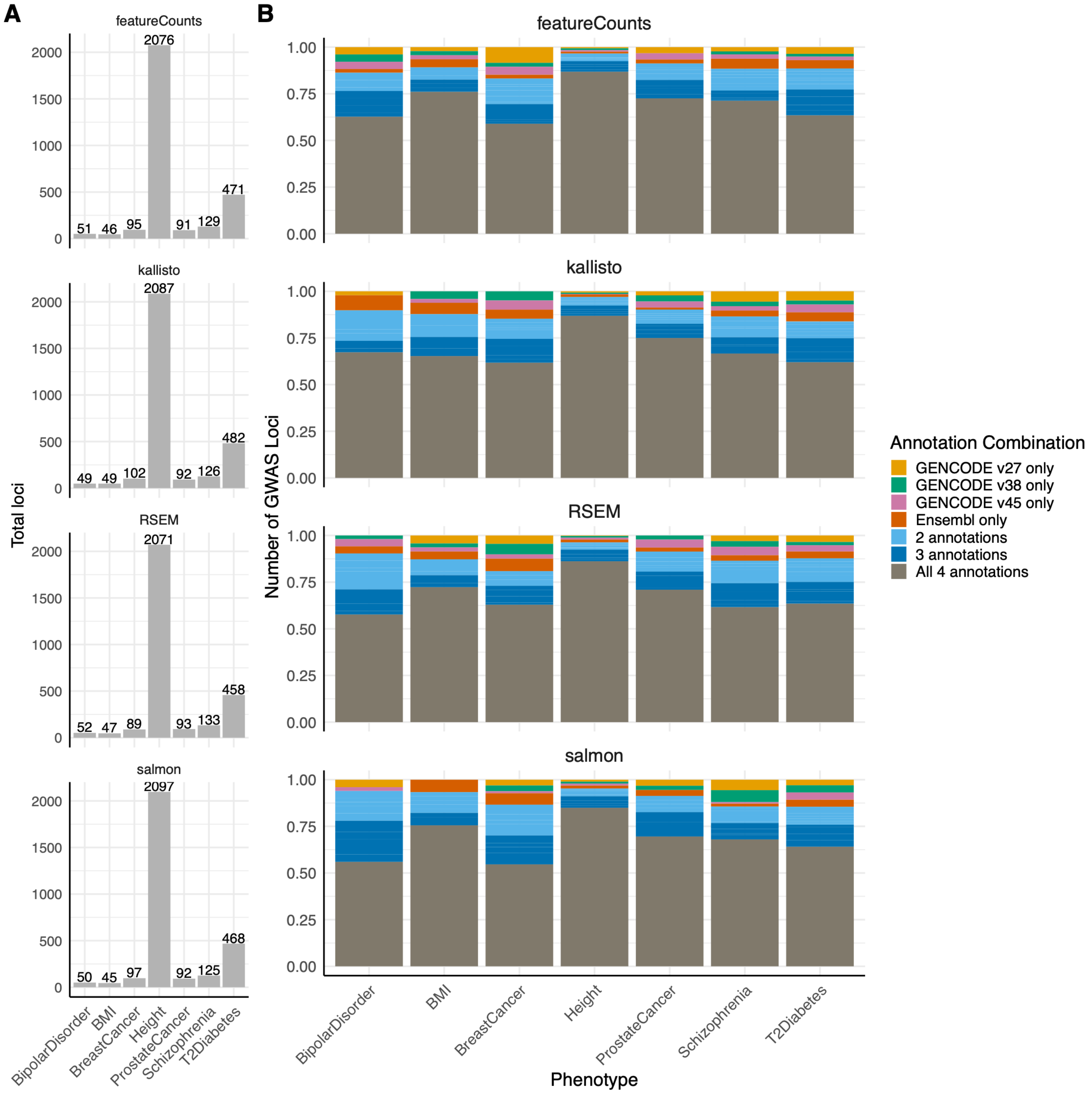


**Figure S28**: **(A)** Total number of GWAS loci tagged (within 1Mb) through colocalization with seven phenotypes across all annotations for a given quantification method. **(B)** The proportion of GWAS loci tagged by different annotation combinations for a given phenotype and quantification method.

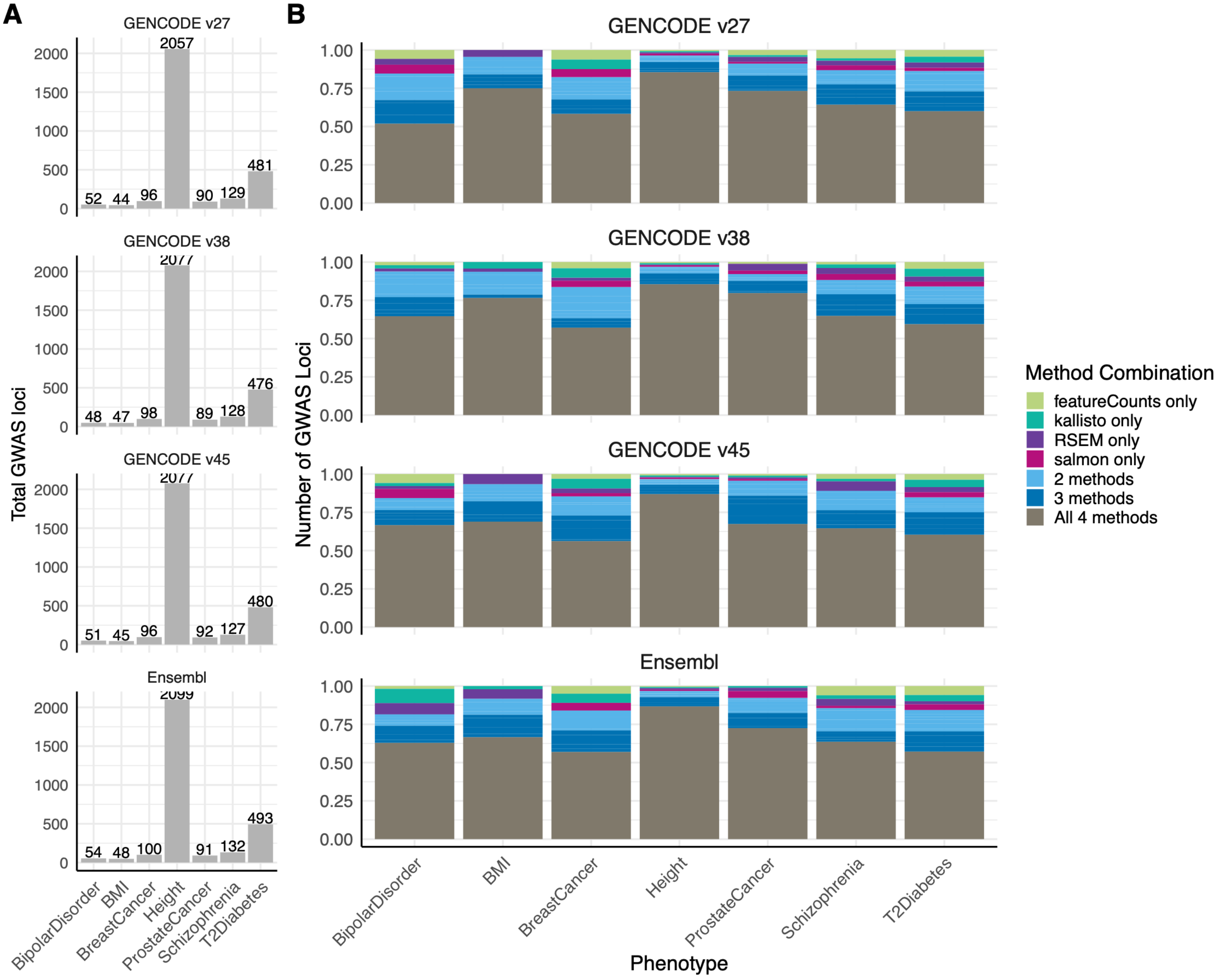


**Figure S29**: **(A)** Total number of GWAS loci tagged (within 1Mb) through colocalization with seven phenotypes across all methods for a given transcriptome annotation. **(B)** The proportion of GWAS loci tagged by different method combinations for a given phenotype and annotation.

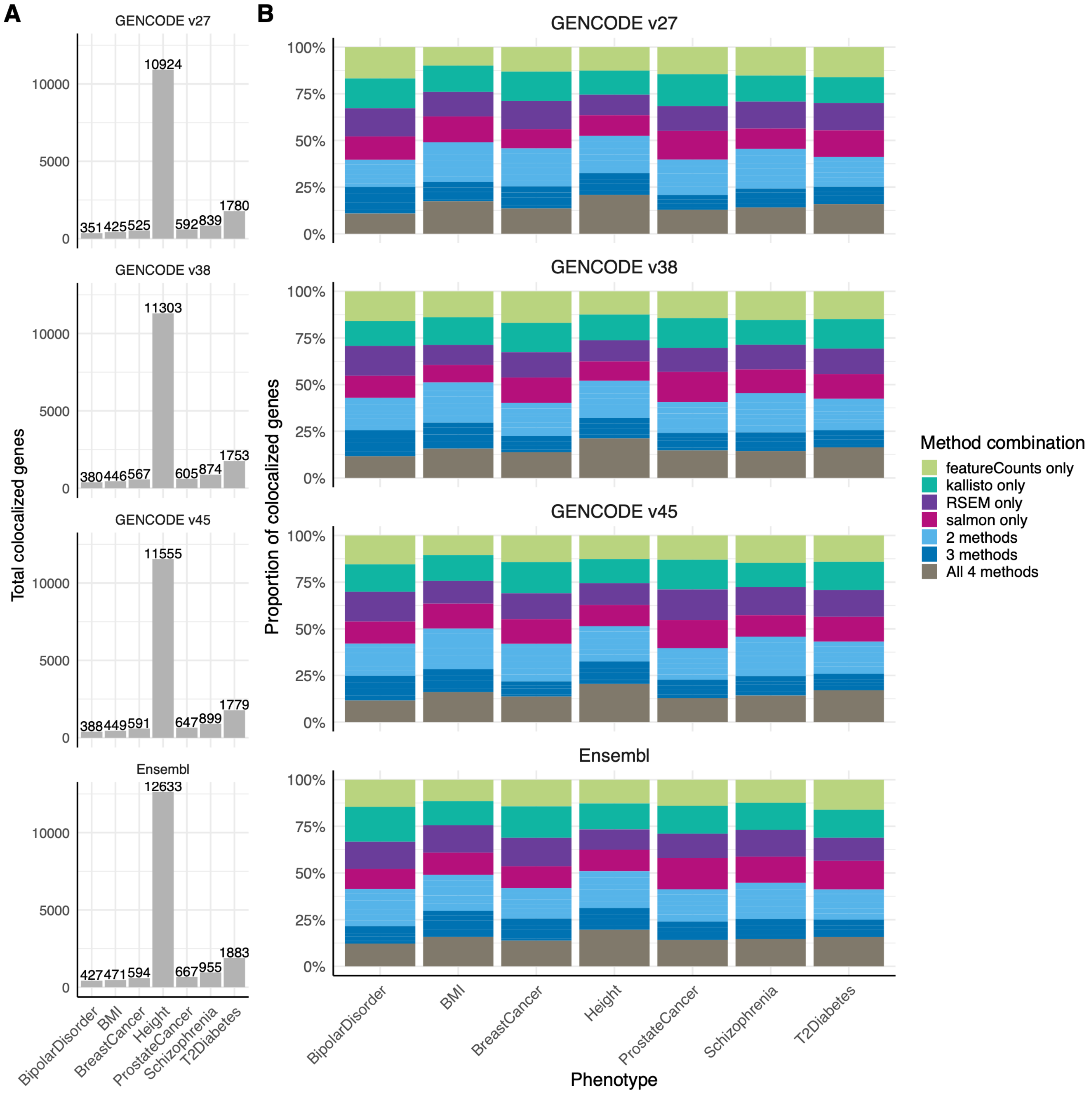


**Figure S30**: **(A)** Total number of genes colocalized at GWAS loci for seven phenotypes across all methods for a given transcriptome annotation. **(B)** The proportion of colocalized genes identified by different method combinations for a given phenotype and annotation.

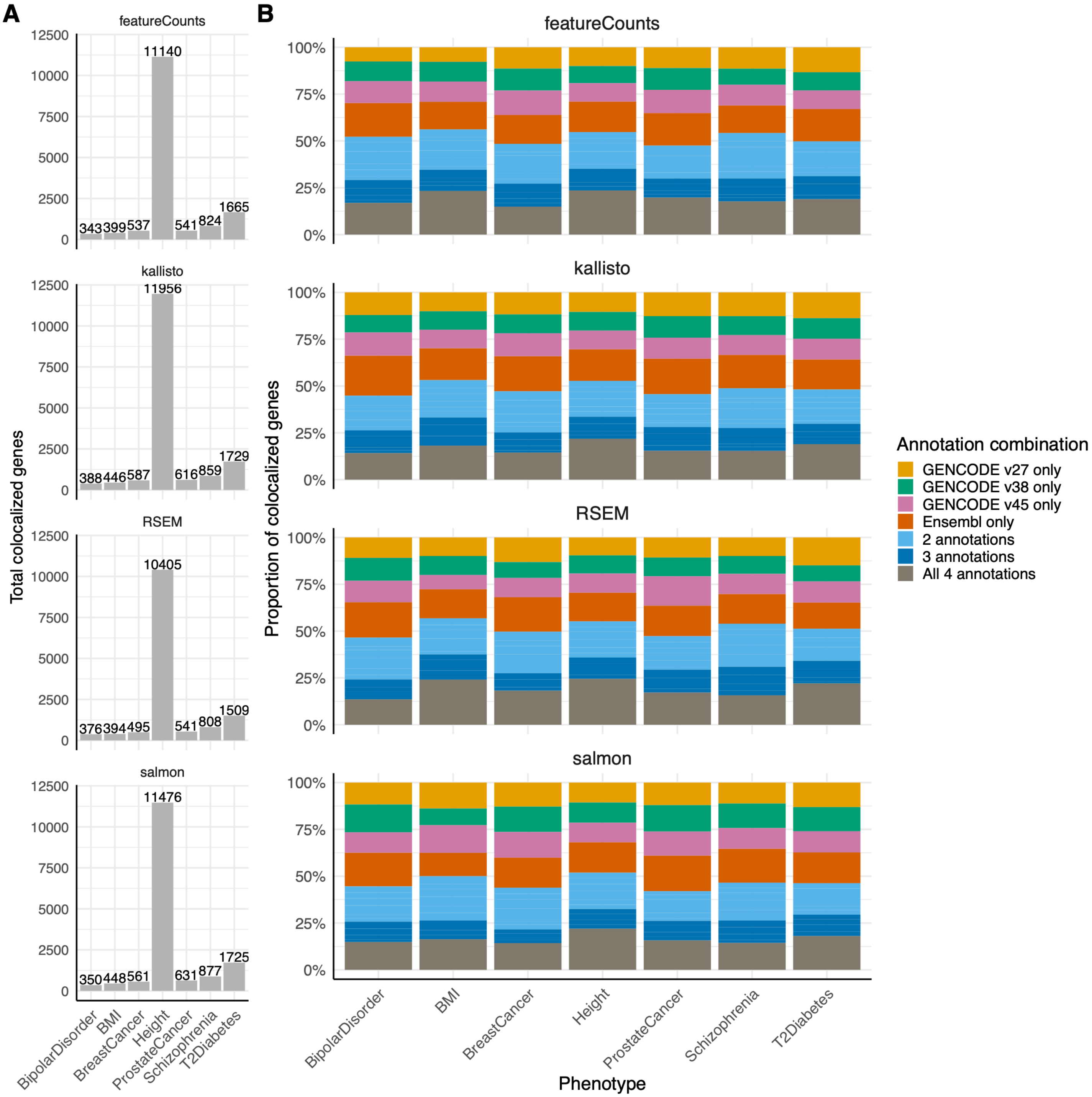


**Figure S31**: **(A)** Total number of genes colocalized at GWAS loci for seven phenotypes across all annotations for a given quantification method. **(B)** The proportion of colocalized genes identified by different annotation combinations for a given phenotype and quantification method.

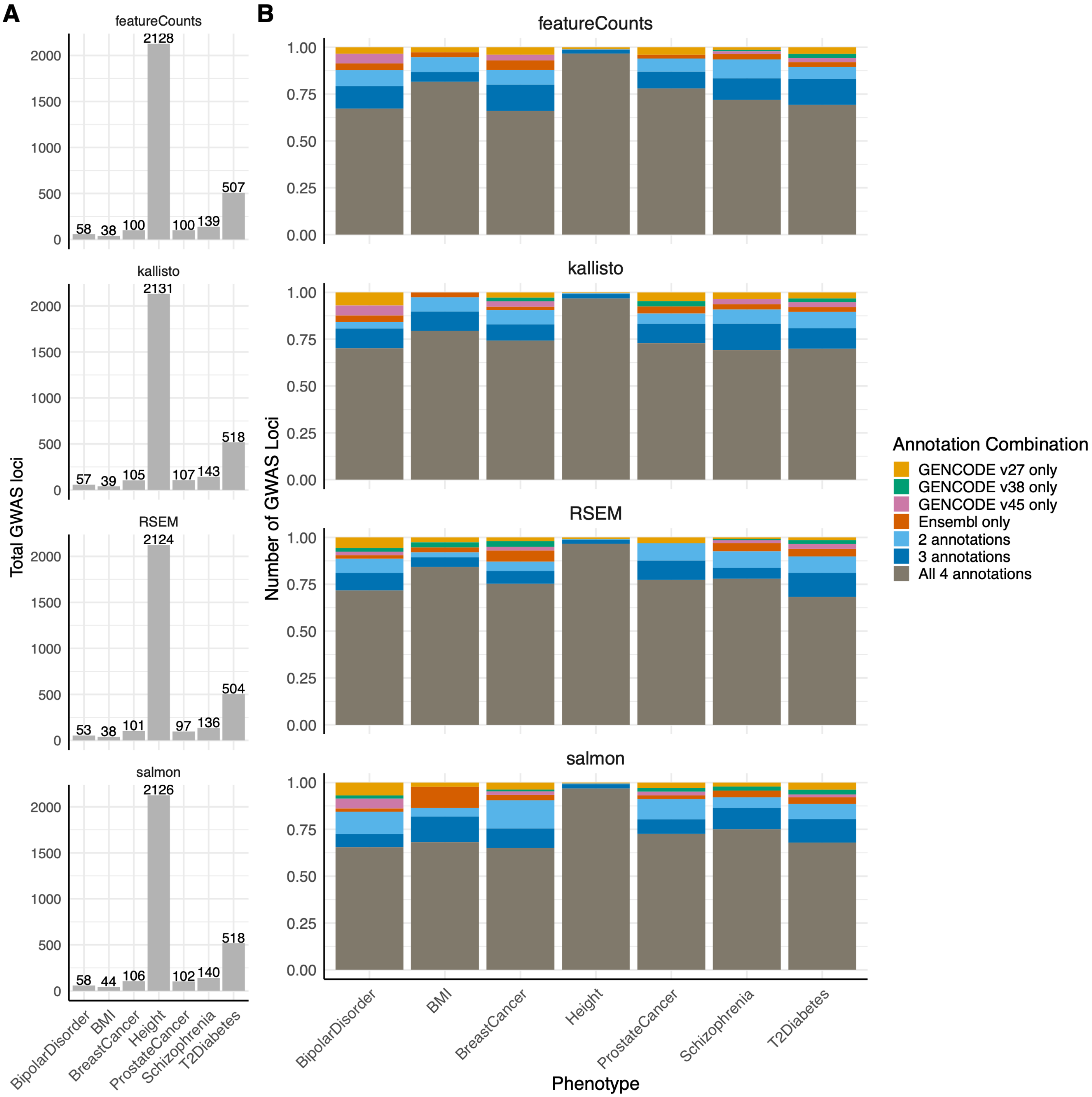


**Figure S32**: **(A)** Total number of GWAS loci tagged (within 1Mb) through TWAS genes with seven phenotypes across all annotations for a given quantification method. **(B)** The proportion of GWAS loci tagged by different annotation combinations for a given phenotype and quantification method.

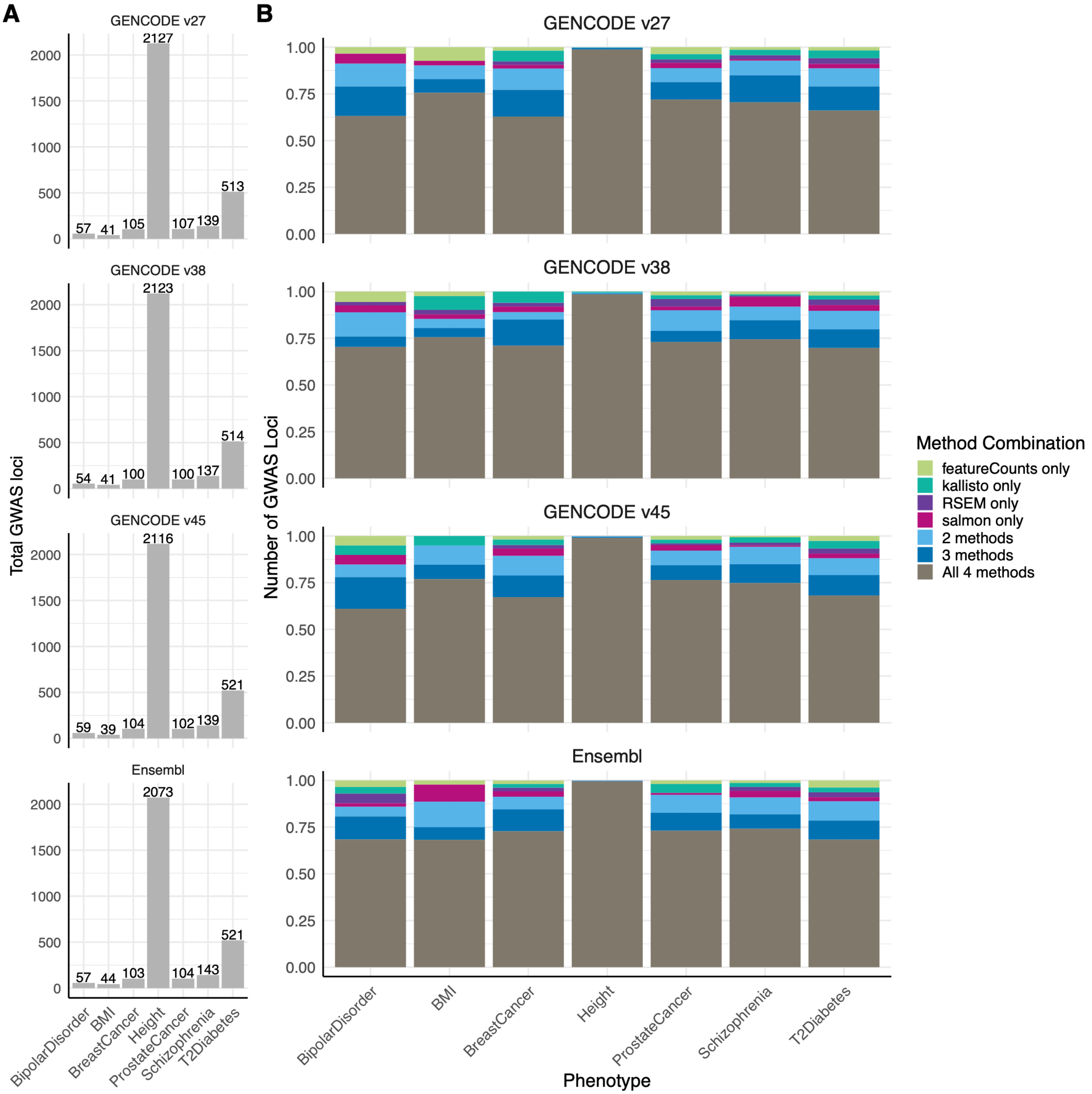


**Figure S33**: **(A)** Total number of GWAS loci tagged (within 1Mb) through TWAS genes with seven phenotypes across all methods for a given transcriptome annotation. **(B)** The proportion of GWAS loci tagged by different method combinations for a given phenotype and annotation.

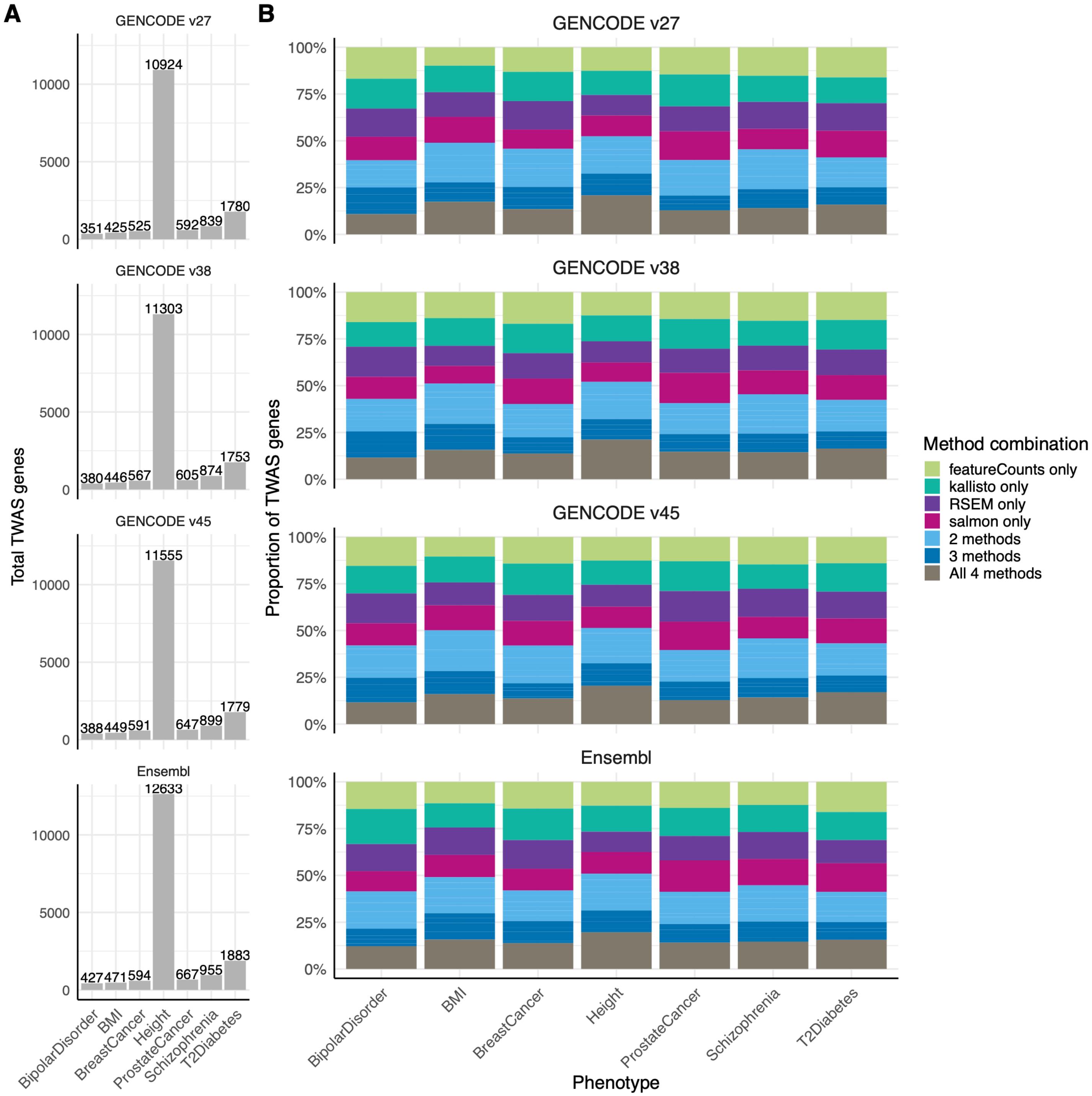


**Figure S34**: **(A)** Total number of TWAS genes for seven phenotypes across all methods for a given transcriptome annotation. **(B)** The proportion of TWAS genes identified by different method combinations for a given phenotype and annotation.

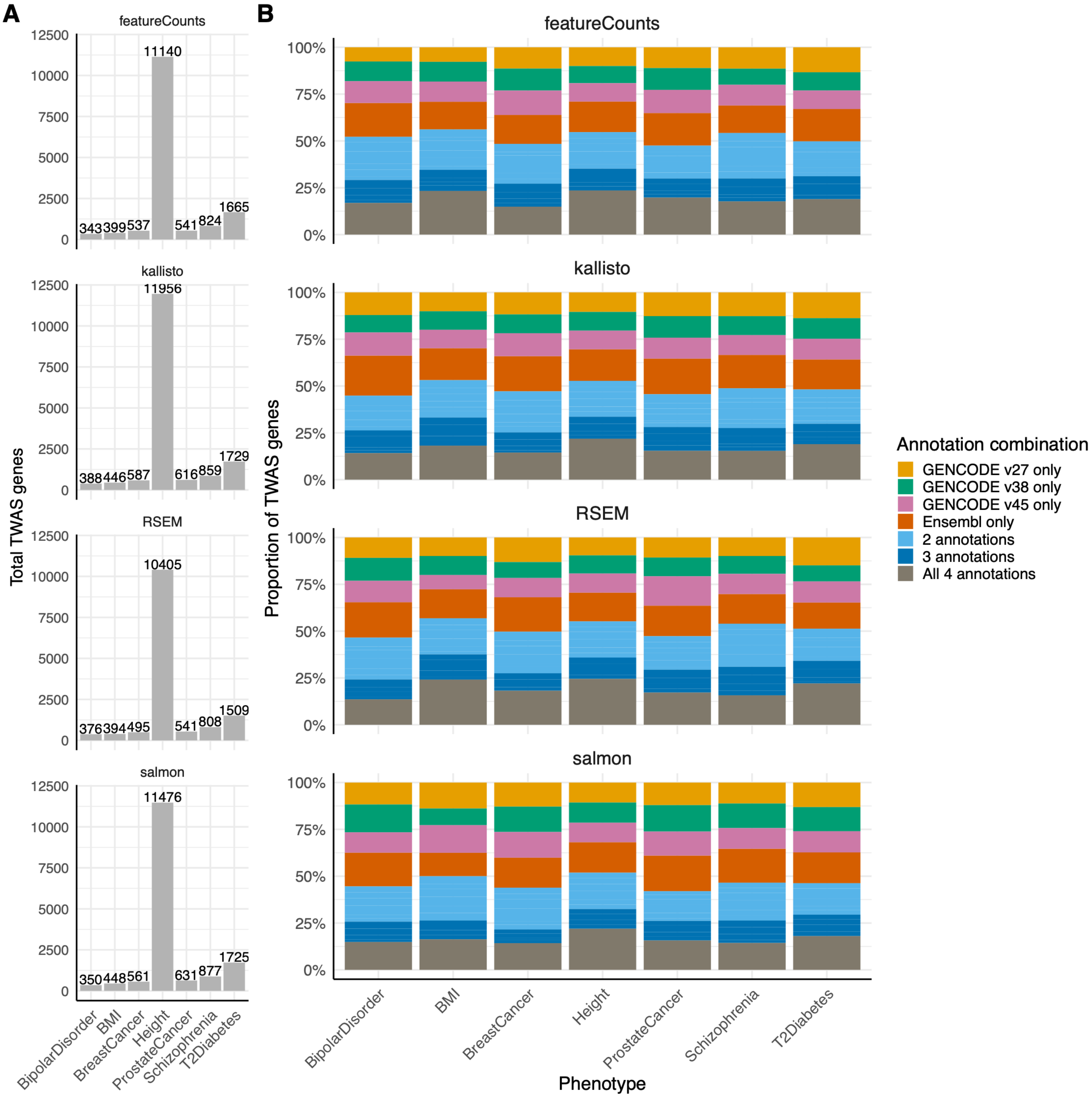


**Figure S35**: **(A)** Total number of TWAS genes for seven phenotypes across all transcriptome annotations for a given quantification method. **(B)** The proportion of TWAS genes identified by different annotation combinations for a given phenotype and method.

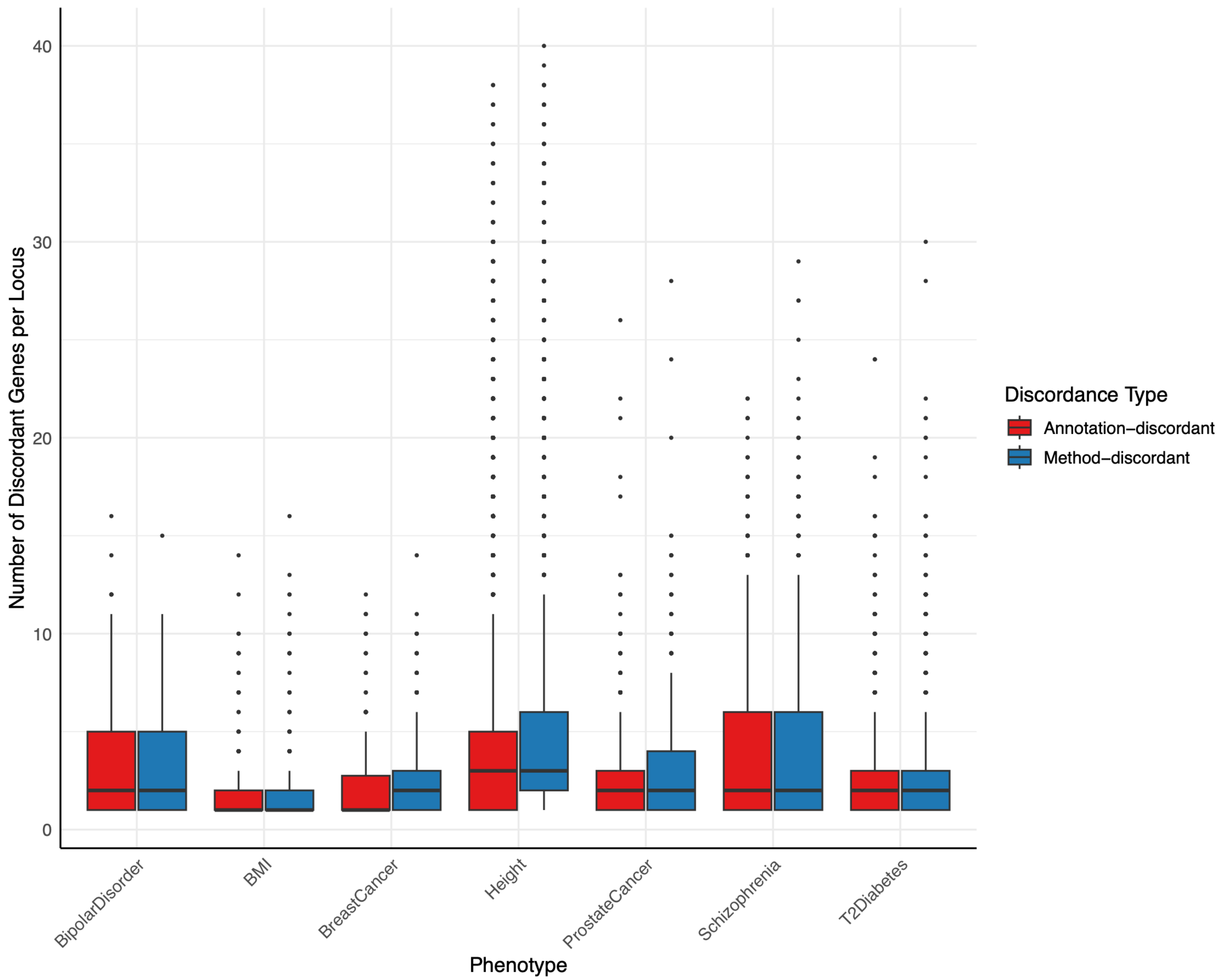


**Figure S36**: Number of discordant genes prioritized by TWAS at the same GWAS locus by discordance type (quantification method [blue], annotation [red]).

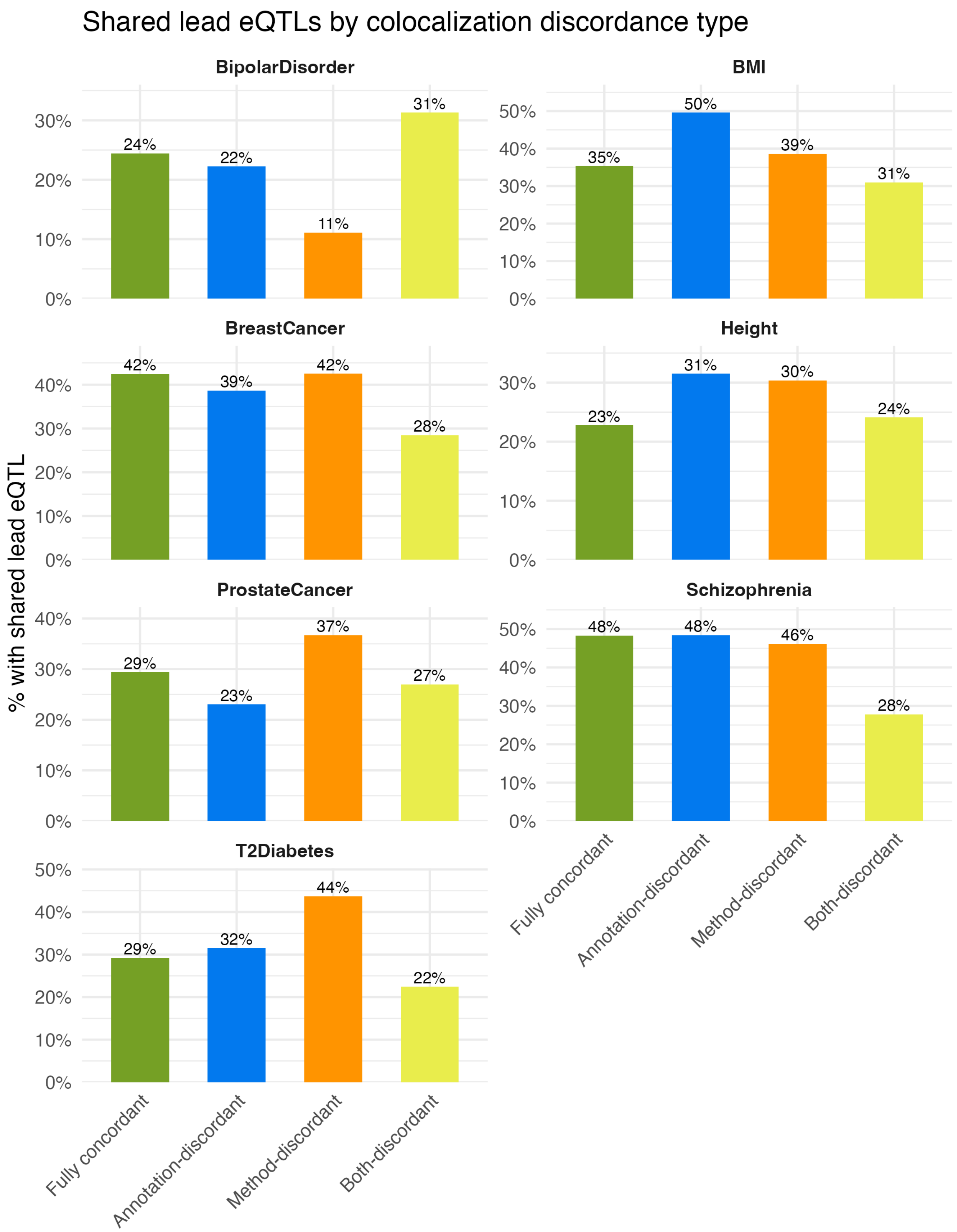


**Figure S37**: Proportion of gene-tissue pairs where the lead eQTL is shared with at least one other gene in the same tissue (assessed across all annotation and method combinations), stratified by colocalization concordance type and faceted by GWAS phenotype. Genes identified in only a single annotation-method setting are excluded.

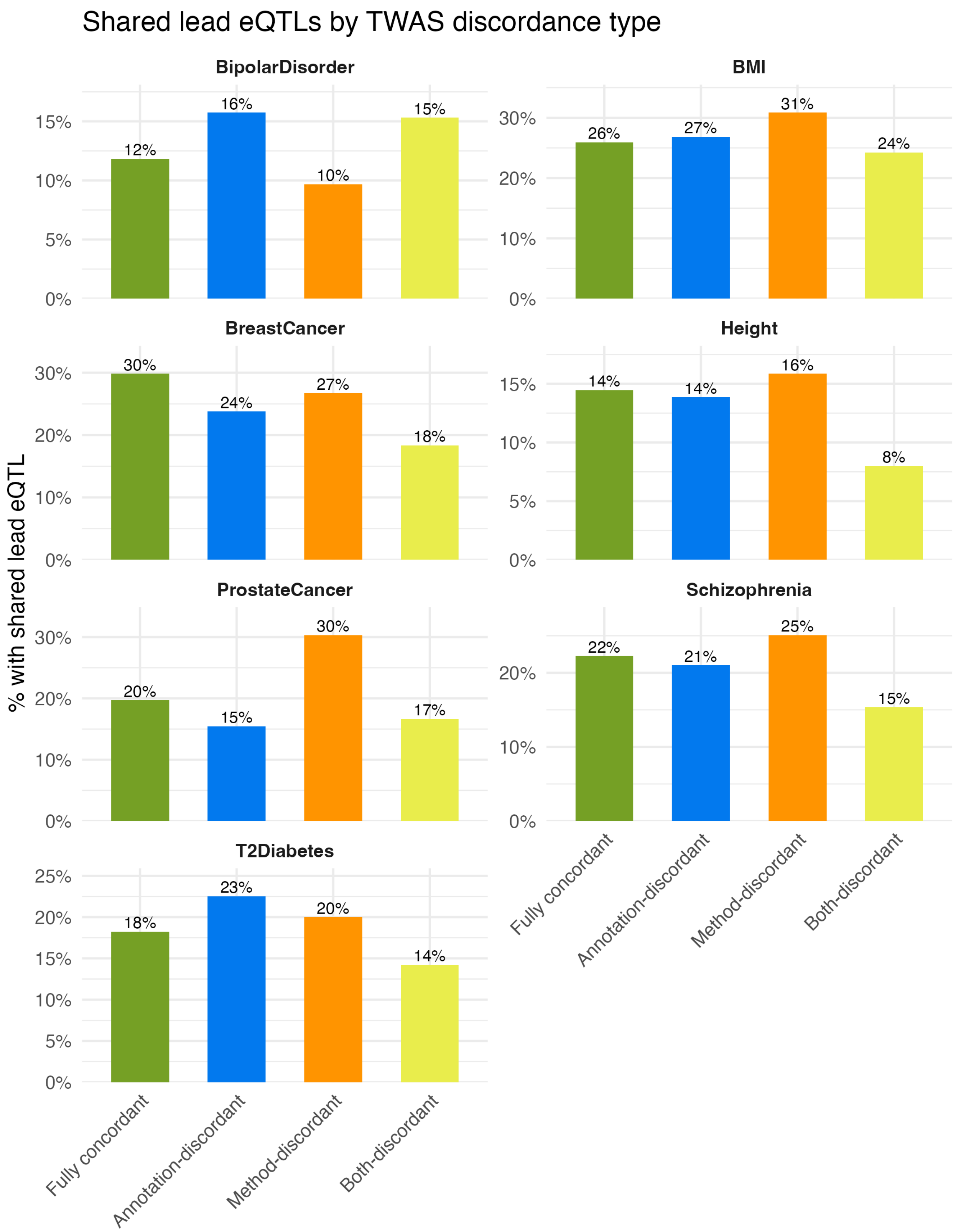


**Figure S38**: Proportion of gene-tissue pairs where the lead eQTL is shared with at least one other gene in the same tissue (assessed across all annotation and method combinations), stratified by TWAS concordance type and faceted by GWAS phenotype. Genes identified in only a single annotation-method setting are excluded.

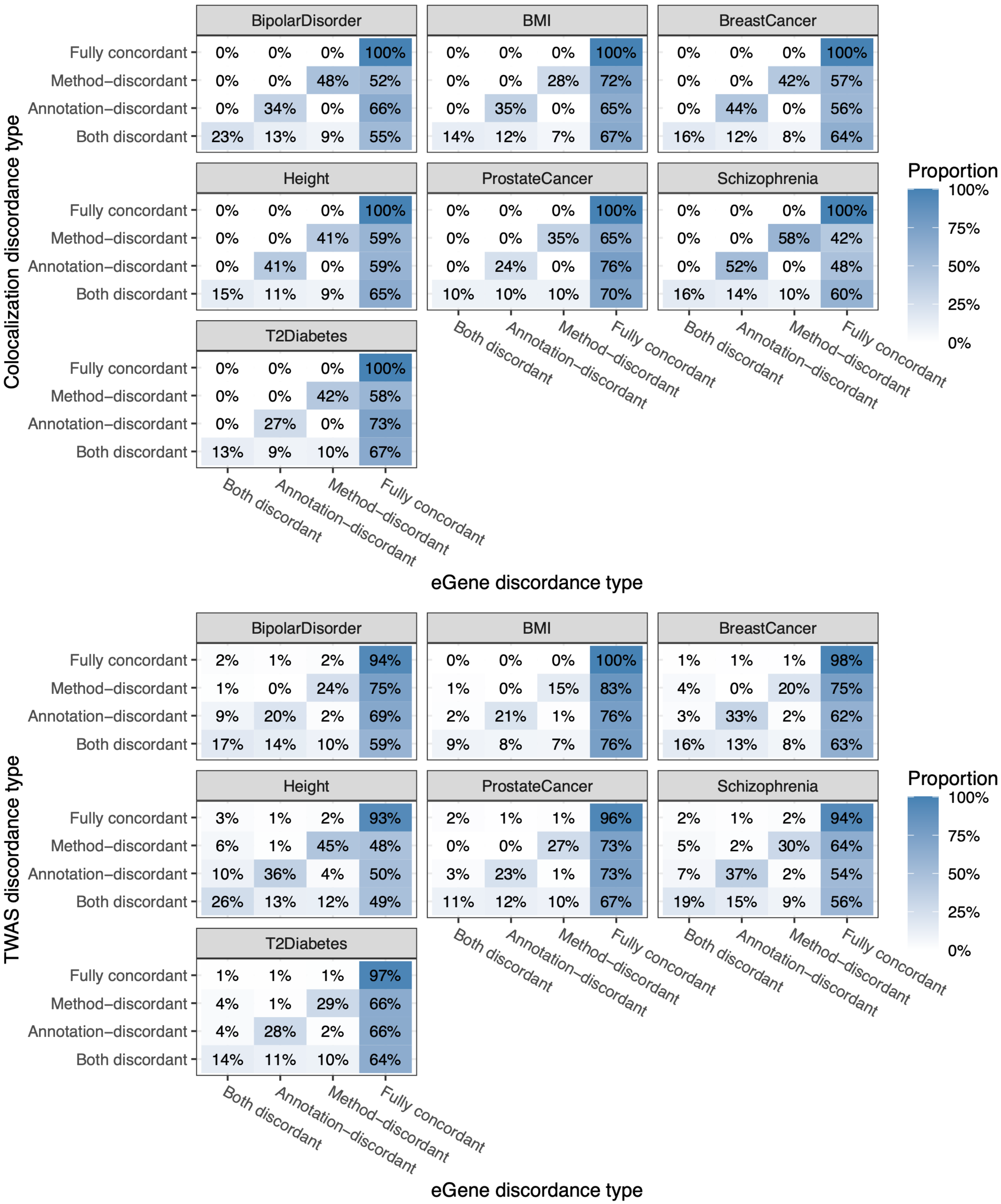


**Figure S39**: The proportion of method-discordant, annotation-discordant, and fully concordant (across both methods and annotations) colocalized genes that are method-discordant, annotation-discordant, or fully concordant eGenes (x-axis; top). The proportion of method-discordant, annotation-discordant, and fully concordant (across both methods and annotations) TWAS-identified genes that are method-discordant, annotation-discordant, or fully concordant eGenes (x-axis; bottom).

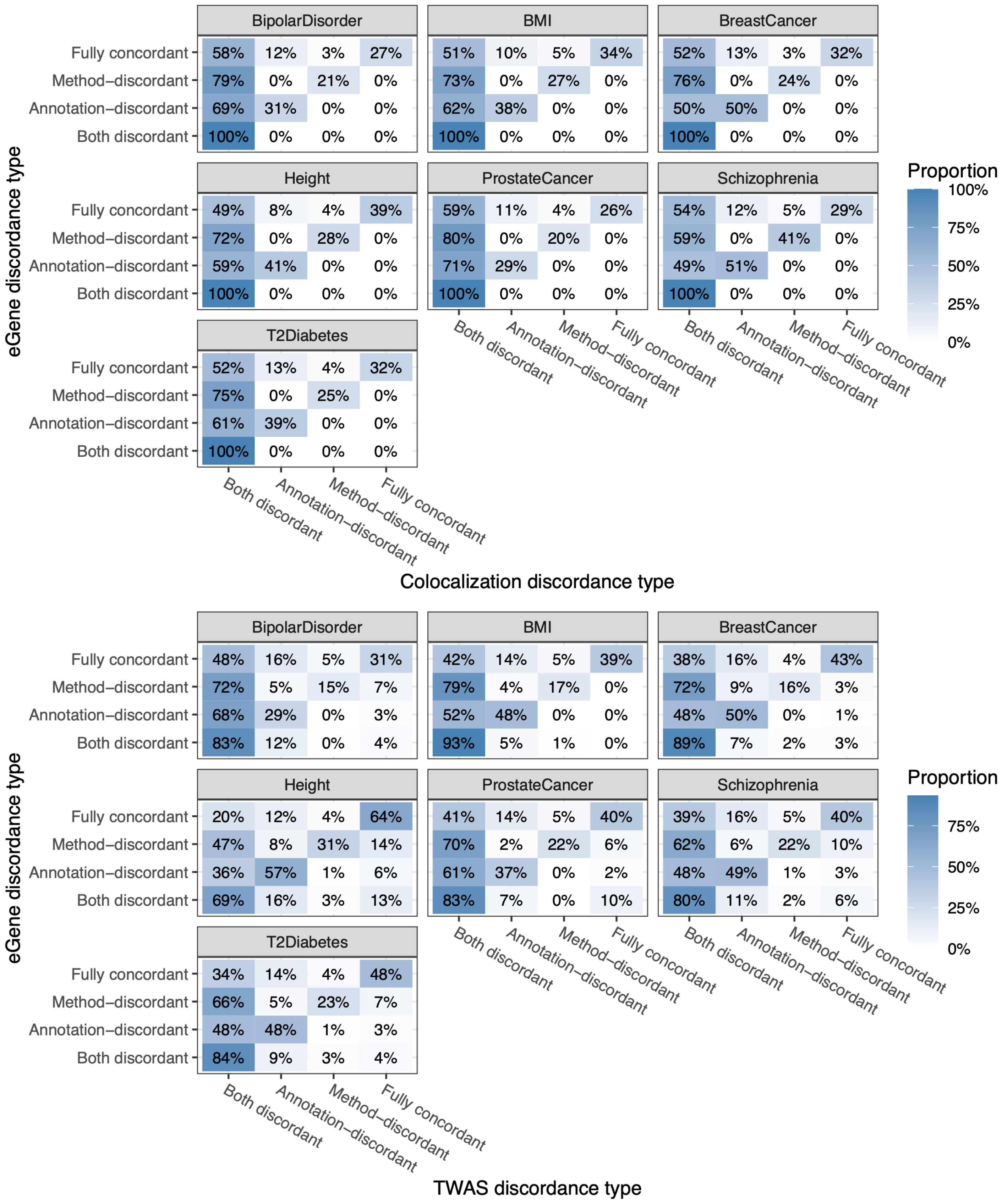


**Figure S40**: The proportion of method-discordant, annotation-discordant, and fully concordant (across both methods and annotations) eGenes that are method-discordant, annotation-discordant, or fully concordant colocalized genes (x-axis; top). The proportion of method-discordant, annotation-discordant, and fully concordant (across both methods and annotations) eGenes that are method-discordant, annotation-discordant, or fully concordant TWAS-identified genes (x-axis; bottom).

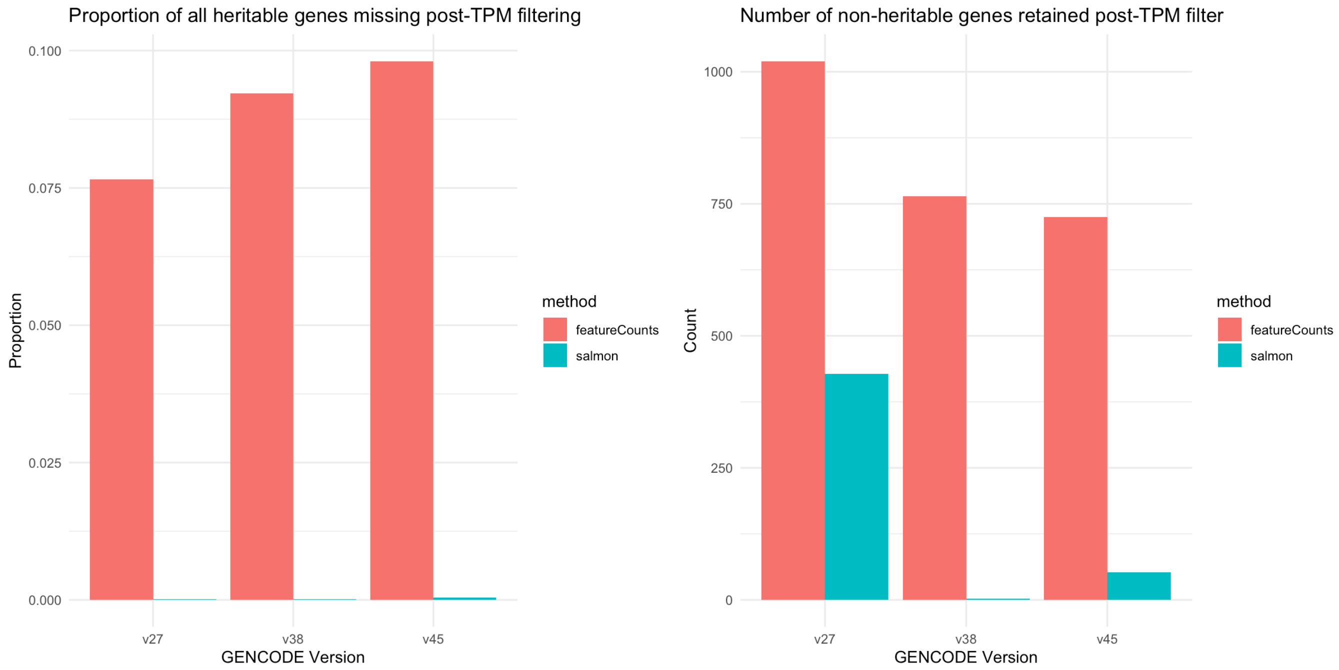


**Figure S41**: (**Left**) Proportion of heritable genes excluded from eGene analysis due to TPM filter by quantification method (STAR+featureCounts [red], Salmon [blue]) for a given annotation. (**Right**) Number of non-genetically-regulated genes retained for eGene analysis by TPM filter by quantification method (STAR+featureCounts [red], Salmon [blue]) for a given annotation.

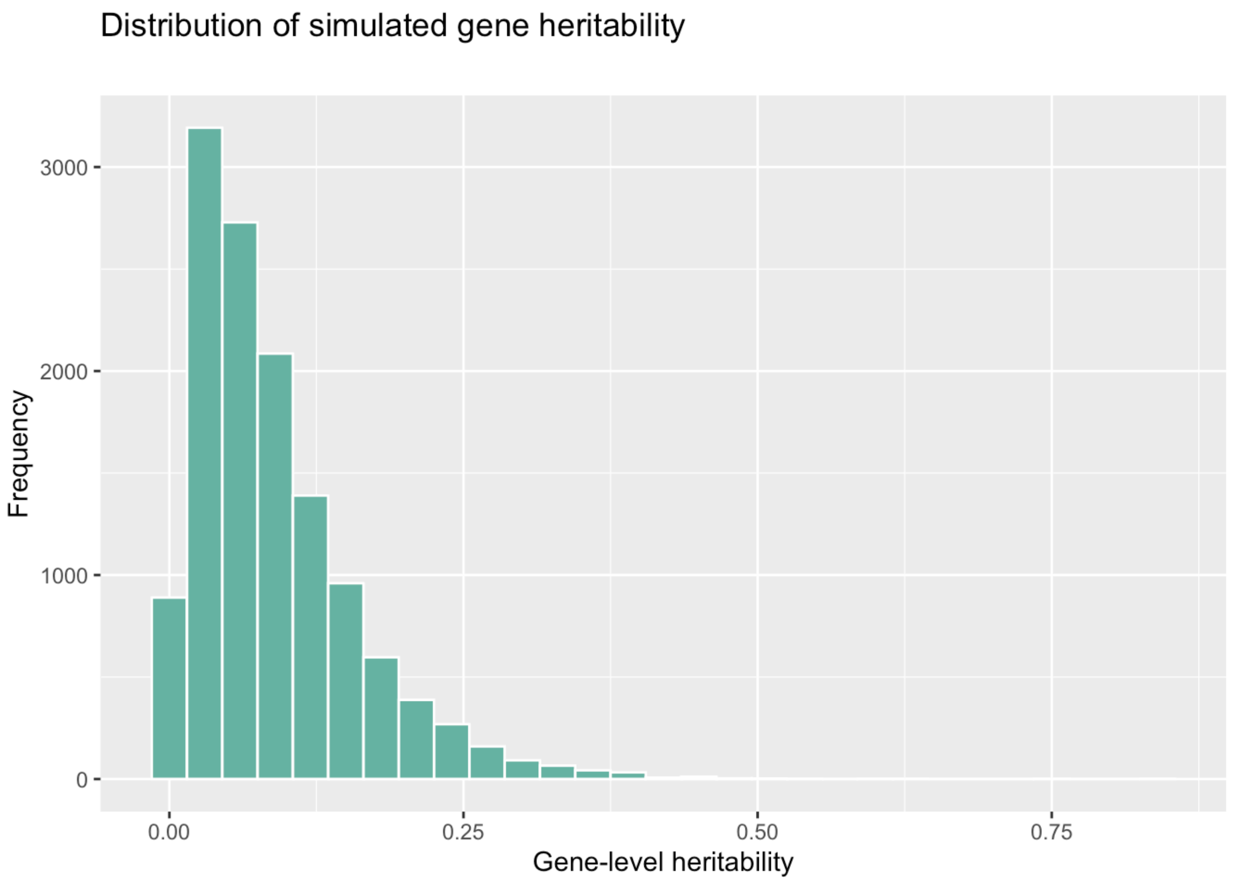

**Figure S42**: Histogram of simulated gene-level heritability across autosomal protein-coding genes present in GENCODE v38.
